# *Cryptococcus neoformans* rewires the conserved Wee1-CDK checkpoint through two divergent kinases required for replication-stress tolerance and virulence

**DOI:** 10.64898/2026.06.24.734400

**Authors:** Jin-Tae Choi, Rodney J. Colón-Reyes, Dong-Hoon Yang, Seung-Heon Lee, Setareh Etesham, Nicole Bodner, Ki Tae Suk, Srikripa Chandrasekaran, Lukasz Kozubowski, Yong-Sun Bahn

**Author notes:** Correspondence and requests for materials should be addressed to LK, and YSB. These authors have contributed equally to this work (Jin-Tae Choi, Rodney J. Colón-Reyes).

## Abstract

Cell-cycle checkpoints couple cell division to environmental and intracellular stress. Here, we show that the human fungal pathogen *Cryptococcus neoformans* possesses two divergent Wee1-family kinases, CnSwe1 and CnSwe102, that retain conserved CDK-inhibitory activity but function differently from their counterparts in the canonical *Saccharomyces cerevisiae* morphogenesis checkpoint. The *swe1*Δ and *swe102*Δ mutants displayed distinct stress-response defects, and genetic analyses suggested a dosage-sensitive genetic interaction between *SWE1* and *SWE102*. Although both proteins promoted Cdc28 tyrosine phosphorylation and elongated-cell morphology when expressed in *S. cerevisiae*, neither localized to the mother-bud neck in *C. neoformans*. Altered *SWE1* dosage in the absence of *SWE102* increased sensitivity to replicative and DNA-damaging stresses and perturbed cell-cycle progression under genotoxic conditions. Importantly, loss of *SWE1* nearly abolished virulence in a murine infection model, and *SWE102* also contributed to pathogenicity. Together, these findings indicate that the conserved Wee1-CDK module acts in *C. neoformans* as a dosage-sensitive checkpoint that promotes stress adaptation and fungal virulence.

## INTRODUCTION

The ability to coordinate cell growth, genome duplication, and cell division is fundamental to fungal proliferation and, consequently, to fungal pathogenesis. During infection, pathogenic fungi encounter host-associated stresses that can perturb DNA replication, cell-wall integrity, cytoskeletal organization, and mitotic progression. In *Cryptococcus neoformans*, cell-cycle-regulated transcriptional programs include genes linked to virulence, and several virulence-associated pathways have been proposed to intersect with cell-cycle progression (Kelliher and Haase, 2017; Kelliher et al., 2016). *C. neoformans* also undergoes extensive physiological and morphological adaptation during dissemination from the lung to the central nervous system, underscoring the importance of coordinated stress adaptation for pathogenic growth (Al-Huthaifi et al., 2024). Cell-cycle checkpoints allow eukaryotic cells to delay cell-cycle transitions under adverse conditions by modulating cyclin-dependent kinase (CDK) activity, thereby preserving genome integrity and cellular viability (Barnum and O’Connell, 2014). Among these surveillance pathways, DNA damage and S-phase checkpoints are particularly relevant to replication-associated stress because they prevent mitotic progression before damaged DNA or stalled replication forks have been properly resolved (Segurado and Tercero, 2009). Thus, cell-cycle control is not merely a housekeeping process in pathogenic fungi, but a potential determinant of stress adaptation and virulence.

Wee1-family kinases are conserved negative regulators of CDKs. In the fission yeast *Schizosaccharomyces pombe*, Wee1 was identified as a dose-dependent inhibitor of mitotic entry; loss of Wee1 activity causes premature division at reduced cell size, whereas increased Wee1 activity delays mitosis (Nurse, 1975; Russell and Nurse, 1987). Mechanistically, Wee1 phosphorylates Cdk1 on a conserved inhibitory tyrosine residue and thereby contributes to G2/M control and DNA damage checkpoint regulation (Raleigh and O’Connell, 2000; Russell and Nurse, 1987). More recent work in other eukaryotic systems has further expanded this view by showing that WEE1 activity can protect stalled replication forks and limit replication-associated genome instability, reinforcing the broader connection between WEE1-dependent CDK control and replication-stress tolerance (Elbaek et al., 2022; Koh, 2022). In the budding yeast *Saccharomyces cerevisiae*, the Wee1 homolog Swe1 phosphorylates Cdc28 and is best known for its role in the morphogenesis checkpoint, which delays nuclear division when bud formation, septin organization, or actin-dependent morphogenesis is perturbed (Lew and Reed, 1995; McMillan et al., 1998; Sia et al., 1996). In this system, Swe1 localization and degradation are tightly linked to the mother-bud neck, where septin-associated factors promote timely Swe1 turnover; defects in this process sustain Cdc28 phosphorylation and produce elongated buds (Longtine et al., 2000; McMillan et al., 1999; McMillan et al., 2002; Sia et al., 1998). However, Wee1-family kinases are not restricted to morphogenesis control, suggesting that the conserved Wee1–CDK module may be wired differently in organisms with distinct cell-cycle architecture and pathogenic lifestyles.

*Cryptococcus neoformans* is a basidiomycetous yeast and a major cause of fungal meningoencephalitis, particularly in immunocompromised individuals. Recent estimates suggest approximately 152,000 HIV-associated cases of cryptococcal meningitis and 112,000 cryptococcal-related deaths annually, highlighting the continuing global burden of cryptococcosis (Rajasingham et al., 2022). Although cryptococcosis remains clinically important, the cell-cycle mechanisms that support cryptococcal survival during infection are still incompletely understood. *C. neoformans* contains a single major CDK, Cdk1, which contributes to bud emergence and DNA synthesis, but its checkpoint regulation is poorly defined (Virtudazo et al., 2010). Notably, the *C. neoformans* genome encodes two predicted Wee1-family kinases, CnSwe1 and CnSwe102. Previous studies have implicated CnSwe1 in survival in human cerebrospinal fluid and CnSwe102 in stress adaptation and virulence, including brain infection-associated phenotypes (Lee et al., 2010; Lee et al., 2020; Lee et al., 2016). These observations suggest that Wee1-family kinases may contribute to cryptococcal pathogenicity, but whether they function as canonical budding-yeast-like morphogenesis checkpoint regulators or instead support a distinct cryptococcal checkpoint program has remained unknown.

Because *C. neoformans* proliferates by budding and forms septin structures at the mother-bud neck, it might be expected to use a septin-dependent, Swe1-mediated morphogenesis checkpoint similar to that of *S. cerevisiae*. Accordingly, cryptococcal cells deficient in the septin complex at the mother-bud neck should exhibit elongated bud morphology due to Swe1-dependent sustained inhibition of Cdk1. However, *C. neoformans* septins have been most clearly linked to morphogenetic events during sexual development and virulence, rather than to a classical elongated-bud arrest phenotype analogous to the *S. cerevisiae* morphogenesis checkpoint under standard vegetative growth conditions (Kozubowski and Heitman, 2010).

Here, we show that *C. neoformans* encodes two divergent but bona fide Wee1-family kinases that retain conserved CDK-inhibitory activity. In contrast to the canonical *S. cerevisiae* morphogenesis checkpoint model, CnSwe1 and CnSwe102 do not concentrate at the mother-bud neck and do not function primarily as bud-neck-associated morphogenesis checkpoint regulators. Instead, our data support a model in which these two kinases function in a partially redundant, dosage-sensitive checkpoint module that promotes tolerance to replication-associated DNA damage and supports cryptococcal virulence. These findings suggest that the Wee1–CDK checkpoint module in *C. neoformans* is distinct from the morphogenesis control described in *S. cerevisiae*, and operates in replication-stress adaptation during pathogenic growth.

## RESULTS

### *C. neoformans* encodes two divergent but structurally related Wee1-family kinases

The *C. neoformans* genome encodes two predicted Wee1-family kinases, CnSwe1 and CnSwe102, encoded by *SWE1* (CNAG_03171) and *SWE102* (CNAG_03369), respectively. To define their evolutionary relationship to known Wee1-family kinases, we performed phylogenetic analysis using full-length amino acid sequences from representative fungal and eukaryotic Wee1-family proteins. This analysis placed CnSwe1 and CnSwe102 as distinct cryptococcal Wee1-family kinases rather than as closely related duplicates within a single branch (Fig. 1A). Comparison with representative species further showed that, unlike the single Swe1/Wee1 kinase found in model yeasts such as *S. cerevisiae* and *S. pombe*, *C. neoformans* contains two Wee1-family homologs, consistent with potential functional diversification of this kinase family in cryptococci.

**Fig. 1.**
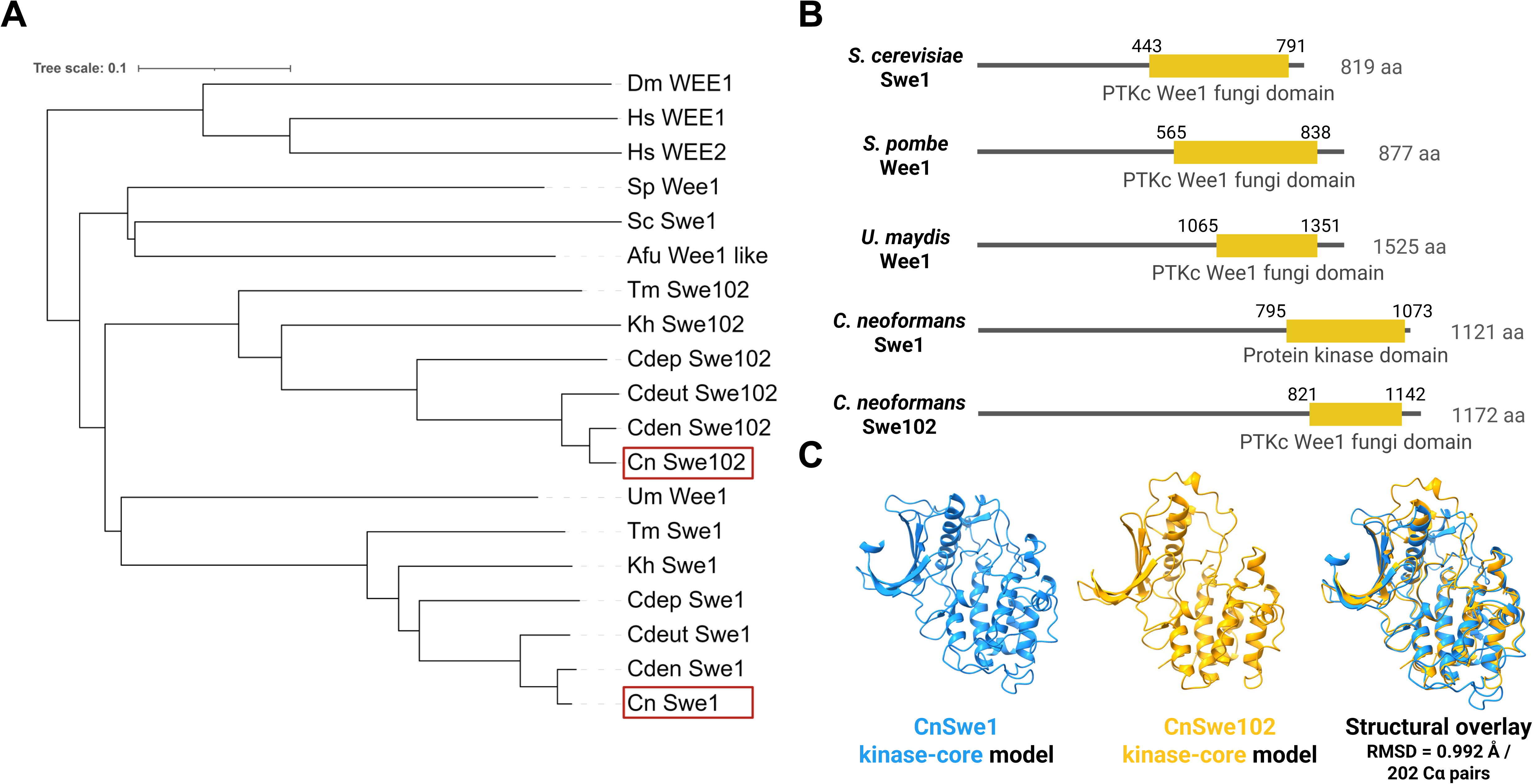
***C. neoformans* encodes two divergent but structurally related Wee1-family kinases.** (A) Phylogenetic analysis of representative Wee1-family kinases. The tree was generated using full-length amino acid sequences of Wee1-family proteins from *Cryptococcus neoformans* H99 (Cn), *Cryptococcus deneoformans* JEC21 (Cden), *Cryptococcus deuterogattii* R265 (Cdeut), *Cryptococcus depauparatus* (Cdep), *Kwoniella heveanensis* (Kh), *Tremella mesenterica* (Tm), *Ustilago maydis* (Um), *Aspergillus fumigatus* (Afu), *Saccharomyces cerevisiae* (Sc), *Schizosaccharomyces pombe* (Sp), *Drosophila melanogaster* (Dm), and *Homo sapiens* (Hs). CnSwe1 and CnSwe102 are highlighted. (B) Domain organization of representative fungal Wee1-family kinases. Protein lengths and predicted C-terminal kinase domains are shown for *S. cerevisiae* Swe1, *S. pombe* Wee1, *U. maydis* Wee1-like kinase, *C. neoformans* CnSwe1, and *C. neoformans* CnSwe102. Domain annotations were based on InterPro analysis. (C) AlphaFold-predicted kinase-core models of CnSwe1 and CnSwe102. The kinase-core regions used for structural comparison correspond to CnSwe1 residues 795–1084 and CnSwe102 residues 818–1146. CnSwe1 is shown in orange and CnSwe102 in blue. Structural superposition was performed using UCSF ChimeraX MatchMaker. RMSD = 0.992 Å over 202 pruned Cα atom pairs.

We next examined the domain organization of representative fungal Wee1-family kinases. InterPro analysis showed that CnSwe1 and CnSwe102 both contain predicted C-terminal Wee1-like kinase domains, despite differences in overall protein length and N-terminal organization (Fig. 1B). This domain architecture resembled the general organization of fungal Wee1-family proteins, in which the conserved catalytic domain is located near the C-terminus. Alignment of conserved kinase-associated motifs further supported the presence of canonical kinase features in both CnSwe1 and CnSwe102, including the ATP-binding/glycine-rich region, Lys-containing kinase subdomain, catalytic HxD-like region, activation-segment motif, and APE/CPE-like motif (Supplementary Fig. 1A).

Because the conservation between CnSwe1 and CnSwe102 was concentrated within the predicted kinase region, we focused the structural comparison on their C-terminal kinase cores. AlphaFold3-predicted kinase-core models of CnSwe1 and CnSwe102 showed a high degree of structural overlap, with an RMSD of 0.992 Å over 202 pruned Cα atom pairs after superposition in UCSF ChimeraX MatchMaker (Fig. 1C). Consistent with this, AlphaFold3 prediction-confidence analysis showed substantially higher mean pLDDT values in the kinase-core regions than across the full-length proteins, supporting the use of the kinase cores for structural comparison (Supplementary Fig. 1B,C). To place these findings in a broader functional context, we compared Wee1-family kinases across representative fungi and mammals. This comparison showed that *C. neoformans* is unusual among the fungal species examined in encoding two Wee1-family kinases, whereas model yeasts such as *S. pombe*, *S. cerevisiae*, and *C. albicans* encode a single Wee1/Swe1 homolog with established roles in CDK inhibitory phosphorylation, cell-size control, or morphogenesis checkpoint regulation (Gale et al., 2009; Lew and Reed, 1995; Nurse, 1975; Russell and Nurse, 1987; Sia et al., 1996) (Table 1). By contrast, mammals encode two Wee1-family paralogs, WEE1/WEE1A and WEE2/WEE1B, with partially specialized roles in CDK inhibition, genome stability, replication control, and oocyte meiosis (Beck et al., 2012; Hanna et al., 2010). Previous mutant-screening and genome-wide kinase studies had suggested that *SWE1* and *SWE102* may be linked to host-relevant survival and virulence-related phenotypes, but the cellular function of CnSwe1 in the H99 background and the functional relationship between the two cryptococcal Wee1-family kinases remained unresolved (Lee et al., 2010; Lee et al., 2020; Lee et al., 2016). Together, these analyses indicate that *C. neoformans* encodes two divergent Wee1-family kinases that differ outside the catalytic region but retain conserved kinase-domain features and structural similarity within their C-terminal kinase cores.

**Table 1.**
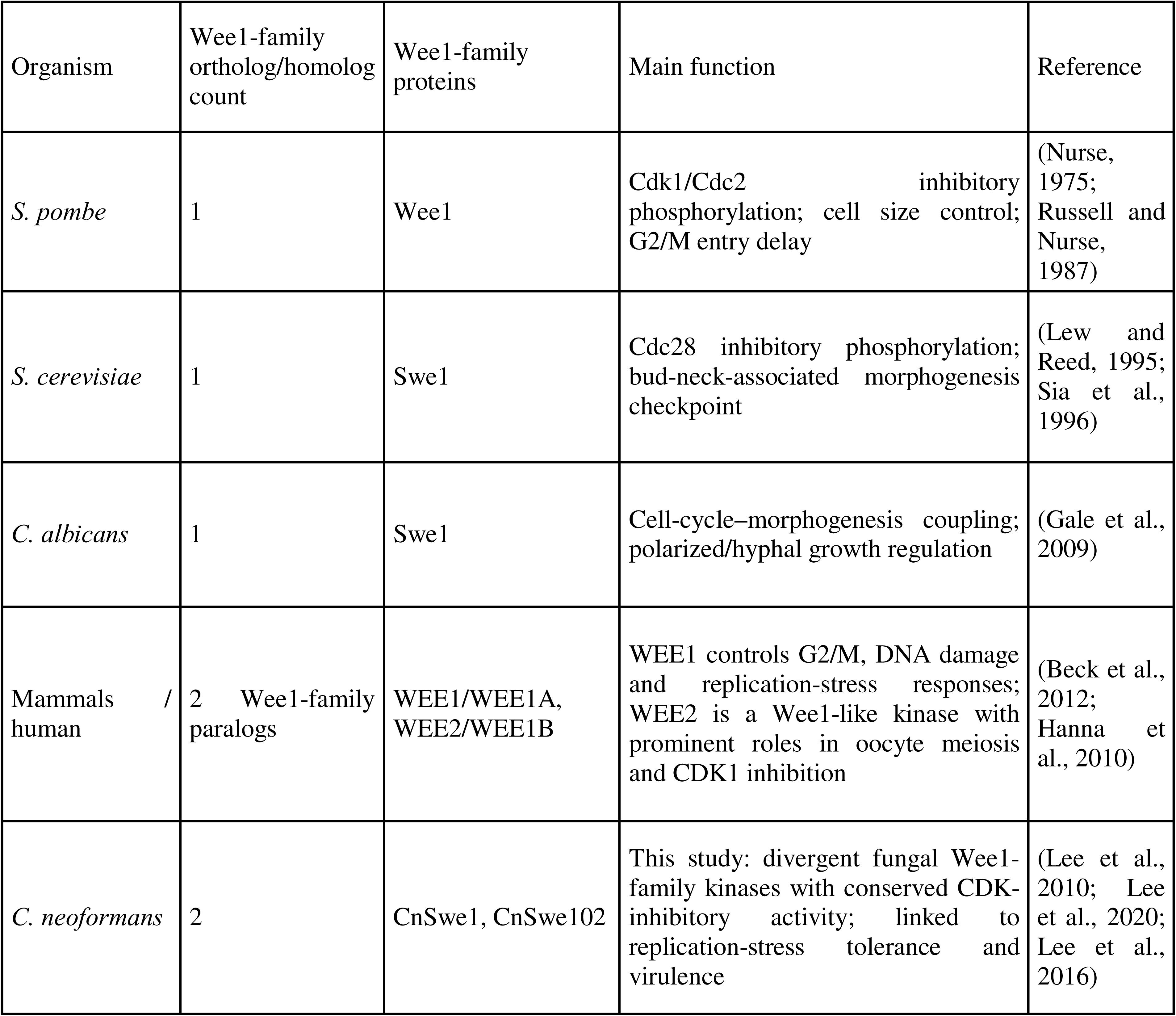
Comparative overview of Wee1-family kinases across representative eukaryotes. Known or proposed Wee1-family kinase functions are summarized for representative fungal and mammalian systems. The functional annotation of CnSwe1 and CnSwe102 reflects the findings of this study.

### CnSwe1 and CnSwe102 mediate overlapping but distinct stress responses in *C. neoformans*

Wee1-family kinases are conserved negative regulators of CDK activity and play central roles in cell-cycle checkpoint control in model yeasts (Lew and Reed, 1995; Russell and Nurse, 1987; Sia et al., 1996). In *C. neoformans*, previous genome-wide functional profiling of kinases implicated CnSwe102 in multiple stress responses, including high temperature, osmotic stress, oxidative stress, DNA damage, ER stress, and fludioxonil susceptibility (Lee et al., 2016). However, the functional contribution of the second cryptococcal Wee1-family kinase, CnSwe1, remained unclear. Whereas the *swe102*Δ mutant was available from our previously constructed genome-wide kinase mutant library (Lee et al., 2016), no deletion mutant of *SWE1* (CNAG_03171) had been generated or phenotypically characterized. We therefore constructed a *swe1*Δ deletion mutant in the *C. neoformans* H99 background by replacing the *SWE1* coding region with a dominant selectable marker (*NAT*; nourseothricin acetyltransferase) through homologous recombination, and confirmed correct gene replacement and the absence of ectopic integration by Southern blot analysis (Supplementary Fig. 2). To verify that the observed mutant phenotypes were attributable to loss of the respective genes, we generated complemented strains by re-integrating a C-terminally tagged *SWE1*-mCherry or *SWE102*-mCherry allele at the corresponding native locus and confirmed correct single-copy integration by diagnostic PCR (Supplementary Fig. 2B).

To investigate the cellular functions of CnSwe1 and CnSwe102, we compared the stress susceptibility profiles of WT, *swe1*Δ, *swe102*Δ, and the corresponding mCherry-tagged complemented strains. The two mutants exhibited both distinct and shared stress-response defects (Fig. 2A). The *swe1*Δ mutant showed pronounced sensitivity under host-associated and ER stress conditions, including growth at 39°C, growth at 37°C with 5% CO, and tunicamycin-induced ER stress, indicating that CnSwe1 plays a major role in adaptation to these conditions. In contrast, the *swe102*Δ mutant displayed stronger defects under diamide-induced oxidative stress and NaCl-induced osmotic stress, suggesting a more prominent role for CnSwe102 in these stress-response pathways. Notably, both mutants exhibited shared susceptibility to fludioxonil, fluconazole, and SDS. Because fludioxonil and SDS are closely associated with membrane and cell integrity stress responses, whereas fluconazole targets ergosterol biosynthesis, these shared defects suggest that CnSwe1 and CnSwe102 converge on cellular processes required for maintaining membrane homeostasis and for adaptation to antifungal stress. Complementation with *SWE1*-mCherry or *SWE102*-mCherry largely restored the corresponding mutant phenotypes, supporting the overall functionality of the tagged alleles (Fig. 2A). However, the *swe102*Δ::*SWE102*-mCherry strain showed a slight residual growth defect at 39°C, suggesting that the mCherry-tagged CnSwe102 protein may have a minor tag-associated or dominant-negative effect under high-temperature stress. Together, these results indicate that CnSwe1 and CnSwe102 perform distinct stress-specific functions while also contributing to a common stress-adaptation network.

**Fig. 2.**
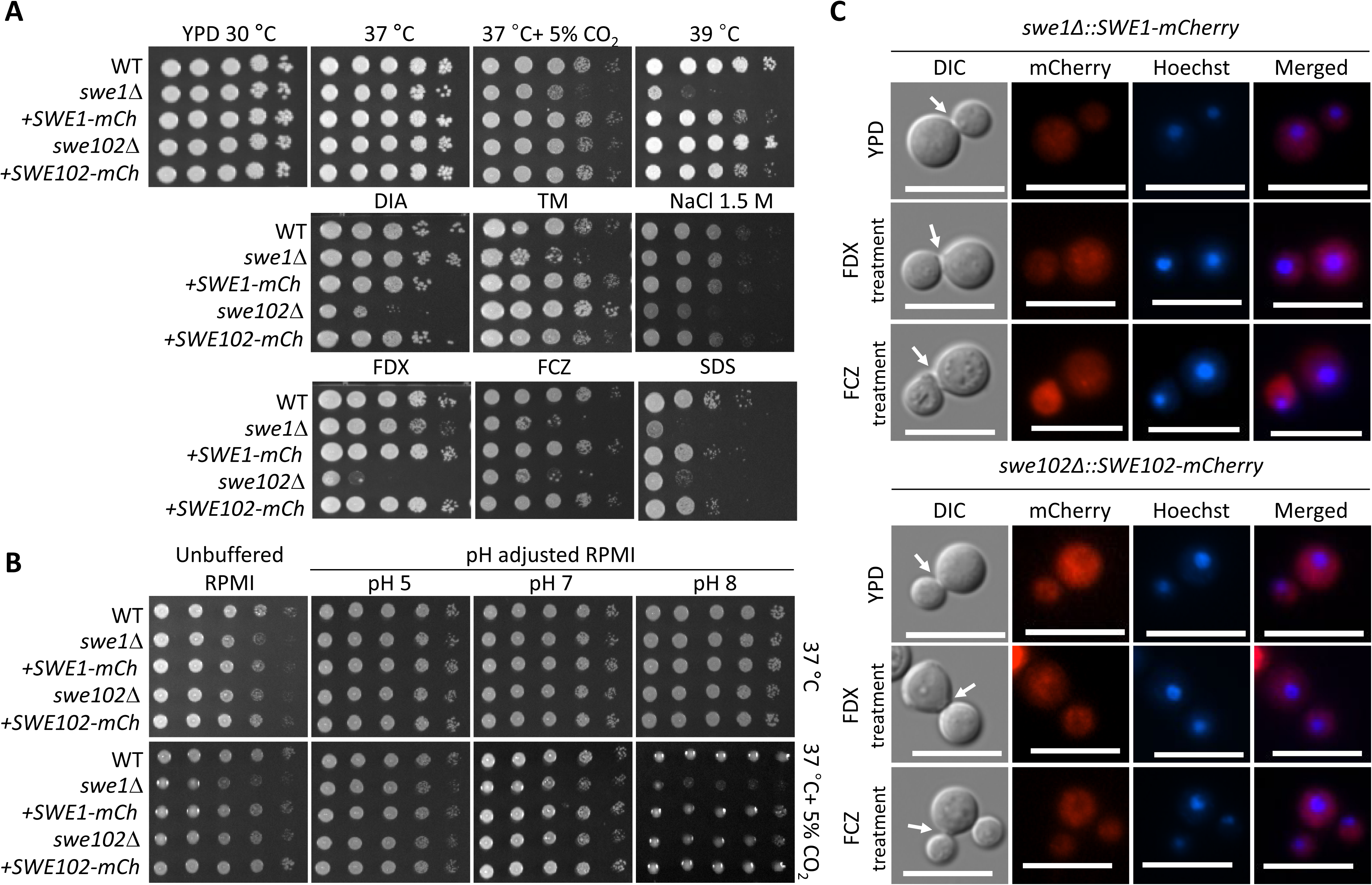
CnSwe1 and CnSwe102 mediate overlapping but distinct stress responses in *C. neoformans*. (A) Serial-dilution growth analysis of WT, *swe1*Δ, *swe102*Δ, *swe1*Δ::*SWE1*-*mCherry*, and *swe102*Δ::*SWE102*-*mCherry* strains under selected stress conditions. Cells were spotted onto YPD agar and incubated under temperature and CO stress conditions or on YPD agar supplemented with the indicated stress-inducing agents: diamide (DIA; 2 mM), tunicamycin (TM; 0.3 μg/ml), NaCl (1.5 M), fludioxonil (FDX; 1 μg/ml), fluconazole (FCZ; 10 μg/ml), or sodium dodecyl sulfate (SDS; 0.03%). (B) Growth of WT, *swe1*Δ, *swe102*Δ, *swe1*Δ::*SWE1*-*mCherry*, and *swe102*Δ::*SWE102*-*mCherry* strains on unbuffered or pH-adjusted RPMI medium. RPMI medium was adjusted to pH 5 (acidic), pH 7 (neutral), or pH 8 (alkaline) as indicated, and plates were incubated at 37 °C in ambient air or at 37 °C with 5% CO . (C) Fluorescence microscopy analysis of CnSwe1-mCherry and CnSwe102-mCherry in the corresponding complemented strains, *swe1*Δ::*SWE1*-*mCherry* and *swe102*Δ::*SWE102*-*mCherry*. Overnight cultures were subcultured into fresh YPD medium at an initial OD_600_ of 0.2 and grown to OD_600_ 0.8. Cells were then left untreated or treated with fludioxonil (FDX; 1 μg/ml) or fluconazole (FCZ; 10 μg/ml) for 1 h. After treatment, cells were collected, stained with Hoechst, and examined by fluorescence microscopy. Differential interference contrast (DIC), mCherry fluorescence, Hoechst-stained nuclei, and merged images are shown. Representative budded cells are shown, and arrows indicate the bud neck.

Because elevated temperature and CO have been linked to pH-dependent adaptation in *C. neoformans* (Chadwick et al., 2022; Choi et al., 2026), we next examined whether CnSwe1 and CnSwe102 contribute to growth under different pH conditions using unbuffered and pH-adjusted RPMI medium. Under most pH-adjusted conditions, the growth defects observed at 37 °C with 5% CO were largely alleviated in both mutants. However, a striking exception was observed under alkaline conditions (pH 8.0) at 37 °C in the presence of 5% CO, where the *swe1*Δ mutant exhibited severe growth sensitivity compared with WT and the complemented strain (Fig. 2B).

These results indicate that CnSwe1 plays a particularly important role in adaptation to alkaline stress when elevated temperature and 5% CO are combined. The complemented strains showed restoration toward WT-like growth, confirming that the observed phenotypes were attributable to loss of the corresponding genes (Fig. 2B). These findings suggest that CnSwe1, in particular, contributes to pH adaptation under host-relevant environmental conditions.

We then assessed the subcellular distribution of CnSwe1 and CnSwe102 using the functional mCherry-tagged complemented strains. After growth in YPD medium or short-term treatment with fludioxonil or fluconazole, CnSwe1-mCherry and CnSwe102-mCherry fluorescence was detected predominantly throughout the cytoplasm, without obvious enrichment at the mother-bud neck under the conditions examined (Fig. 2C). Together with the stress susceptibility data, these observations suggest that CnSwe1 and CnSwe102 function in overlapping but distinct stress-response pathways in *C. neoformans*, while their localization pattern differs from the canonical bud-neck-associated Swe1 organization described in *S. cerevisiae* (Longtine et al., 2000; McMillan et al., 1999).

### Functional interplay between CnSwe1 and CnSwe102 is required for viability, stress adaptation, and morphogenesis

The partially overlapping stress-response phenotypes of the *swe1*Δ and *swe102*Δ mutants suggested that CnSwe1 and CnSwe102 function in related pathways, whereas their distinct phenotypic differences indicated that the two kinases are not functionally equivalent (Fig. 2). This combination of shared and distinct phenotypes suggested that the two cryptococcal Wee1-family kinases may provide partially compensatory but non-identical functions. To further examine their genetic relationship, we attempted to construct a *swe1*Δ *swe102*Δ double mutant.

However, repeated attempts to obtain this strain were unsuccessful, suggesting that simultaneous loss of both kinases is lethal or severely deleterious (i.e., a synthetic lethal or synthetic sick relationship between *SWE1* and *SWE102*). We therefore used an alternative approach in which *SWE1* expression was conditionally regulated in the *swe102*Δ background. To generate this strain, the native *SWE1* promoter was replaced with the copper-responsive *CTR4* promoter. In this system, CuSO_4_ represses P*_CTR4_*:*SWE1* expression, whereas bathocuproinedisulfonic acid (BCS), a copper chelator, induces expression (Fig. 3A). qRT–PCR analysis confirmed that the *CTR4* promoter enabled tight regulation of *SWE1* expression in the *swe102*Δ background: CuSO_4_ strongly reduced *SWE1* transcript levels, whereas BCS increased *SWE1* expression by approximately 2.5-fold relative to the untreated condition (Fig. 3B). Consistent with this transcriptional regulation, growth assays under the same copper-repressive and BCS-inducing conditions showed corresponding growth responses, further validating that the P*_CTR4_*:*SWE1* system tightly controls *SWE1* function at the phenotypic level (Fig. 3C; Supplementary Fig. 4).

**Fig. 3.**
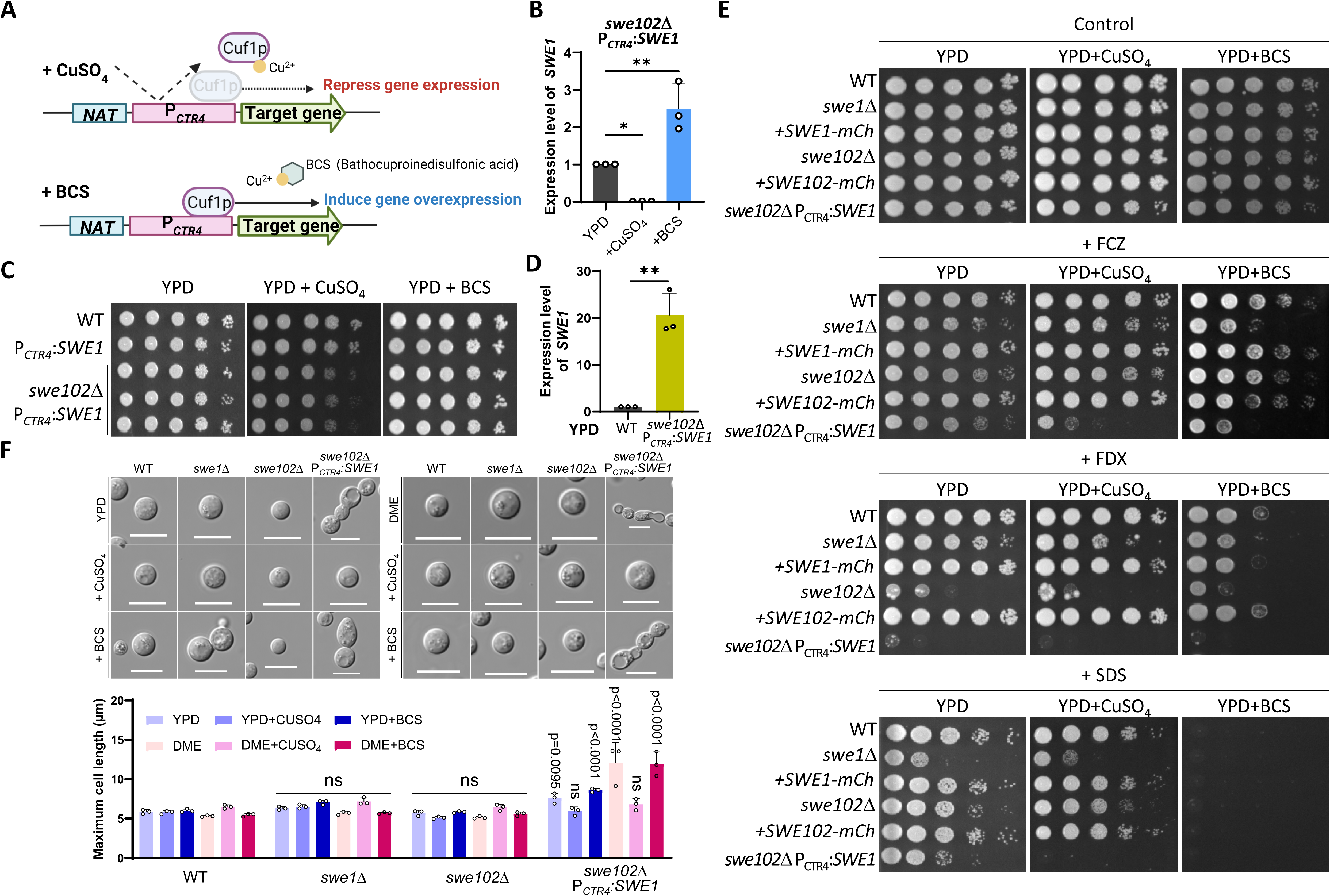
Conditional regulation of *SWE1* in the *swe102*Δ background reveals dosage-dependent effects on growth, morphology and stress susceptibility. (A) Schematic representation of the copper-responsive *CTR4* promoter replacement system used to conditionally regulate target gene expression. Addition of CuSO_4_ represses *PCTR4*-driven expression, whereas bathocuproinedisulfonic acid (BCS), a copper chelator, induces gene overexpression. (B) qRT–PCR analysis of *SWE1* transcript levels in the *swe102*Δ P*_CTR4_:SWE1* strain grown in YPD, YPD + CuSO_4_ (25 μM), or YPD + BCS (200 μM) medium. Transcript levels were normalized to *ACT1*. Data are shown as mean ± SD. Statistical significance was determined by ordinary one-way ANOVA followed by Tukey’s multiple-comparisons test. (C) Serial-dilution growth analysis of WT, P*_CTR4_:SWE1* and *swe102*Δ P*_CTR4_:SWE1* strains on YPD, YPD + CuSO_4_ and YPD + BCS media. (D) qRT–PCR analysis of *SWE1* transcript levels in WT and *swe102*Δ P*_CTR4_:SWE1* strains grown in YPD medium. Transcript levels were normalized to *ACT1*. Data are shown as mean ± SD. Statistical significance was determined by an unpaired two-tailed Student’s t-test. (E) Serial-dilution growth analysis of WT, *swe1*Δ, *swe102*Δ, *swe1*Δ*::SWE1-mCherry*, *swe102*Δ*::SWE102-mCherry* and *swe102*Δ P*_CTR4_:SWE1* strains under basal and stress conditions. Cells were spotted onto YPD, YPD + CuSO_4_ or YPD + BCS medium without additional stress or supplemented with fluconazole (FCZ; 10 μg/ml), fludioxonil (FDX; 1 μg/ml) or sodium dodecyl sulfate (SDS; 0.03%). (F) Representative differential interference contrast (DIC) microscopy images and quantification of cell length for WT, *swe1*Δ, *swe102*Δ, and *swe102*Δ P*_CTR4_:SWE1* strains grown on YPD or DME medium with or without CuSO_4_ or BCS. For round cells, the cell diameter was measured; for elongated or unseparated cells, the maximum cell length was measured. Scale bars, 10 μm. Statistical significance was determined by ordinary two-way ANOVA followed by Tukey’s multiple-comparisons test.

Notably, even under standard YPD conditions without CuSO_4_ or BCS treatment, *SWE1* transcript levels were markedly elevated in the *swe102*Δ P*_CTR4_*:*SWE1* strain compared with WT (Fig. 3D). Therefore, under basal YPD conditions, this strain exists in a state of elevated *SWE1* expression in the absence of *SWE102*, providing a useful genetic background to assess the consequences of altered *SWE1* dosage. We therefore used this strain to examine how altered *SWE1* dosage in the *swe102*Δ background affects stress adaptation. In an extended stress screen, the *swe102*Δ P*_CTR4_*:*SWE1* strain showed increased susceptibility to several stresses, including fludioxonil, fluconazole, hydroxyurea, methyl methanesulfonate, SDS, CdSO_4_, and NaCl (Supplementary Fig. 3). Among these, fludioxonil, fluconazole, and SDS were also conditions in which both the *swe1*Δ and *swe102*Δ single mutants also showed stress-response defects, suggesting that proper dosage balance between CnSwe1 and CnSwe102 is important for shared antifungal and membrane-associated stress responses.

To test this dosage relationship more directly under shared stress conditions, we examined growth after CuSO_4_-mediated repression or BCS-mediated induction of *SWE1* in the *swe102*Δ P*_CTR4_*:*SWE1* strain. Under fluconazole, fludioxonil, and SDS stress, altered *SWE1* regulation in the *swe102*Δ background produced distinct growth responses compared with WT and the single-mutant strains (Fig. 3E). These results indicate that stress adaptation in the absence of *SWE102* is sensitive to *SWE1* dosage.

Interestingly, we found that modulation of *SWE1* expression in the *swe102*Δ background resulted in pronounced morphological alterations. The *swe102*Δ P*_CTR4_*:*SWE1* strain produced elongated cells and cells with incomplete separation defects under specific conditions, particularly on DME medium and under BCS-induced *SWE1* expression conditions (Fig. 3F). Quantitative analysis of maximum cell length confirmed a significant increase in cell length under these conditions compared with the wild-type strain (Fig. 3F). Together, these results indicate that CnSwe1 and CnSwe102 function in a dosage-sensitive, mutually dependent relationship consistent with synthetic lethality or synthetic sickness when both activities are lost and that perturbation of this balance affects stress susceptibility and cellular morphogenesis.

### CnSwe1 and CnSwe102 retain conserved Wee1-like CDK inhibitory activity

In *S. cerevisiae*, Swe1 delays mitotic entry by phosphorylating Cdc28 and is regulated through Hsl1/Hsl7- and septin-associated mechanisms at the mother-bud neck (Longtine et al., 2000; McMillan et al., 1999; Sia et al., 1996). Because CnSwe1 and CnSwe102 did not show obvious mother-bud-neck enrichment in *C. neoformans* (Fig. 2C), we asked whether these proteins nevertheless retain the conserved biochemical function of Wee1-family kinases: inhibitory phosphorylation of CDK.

The dosage-sensitive growth and morphology phenotypes observed in the *swe102*Δ P*_CTR4_*:*SWE1* strain suggested that CnSwe1 and CnSwe102 may influence cell-cycle control through the conserved Wee1–CDK axis. We first examined whether deletion of either kinase altered inhibitory tyrosine phosphorylation of cryptococcal CDK. Immunoblot analysis using an anti-phospho-Cdc2 Tyr15 antibody showed that CDK phosphorylation remained detectable in both the *swe1*Δ and *swe102*Δ mutants, whereas anti-PSTAIR detection confirmed comparable total CDK levels (Fig. 4A). This result is consistent with the possibility that either kinase alone is sufficient to maintain detectable CDK tyrosine phosphorylation in *C. neoformans*.

**Fig. 4.**
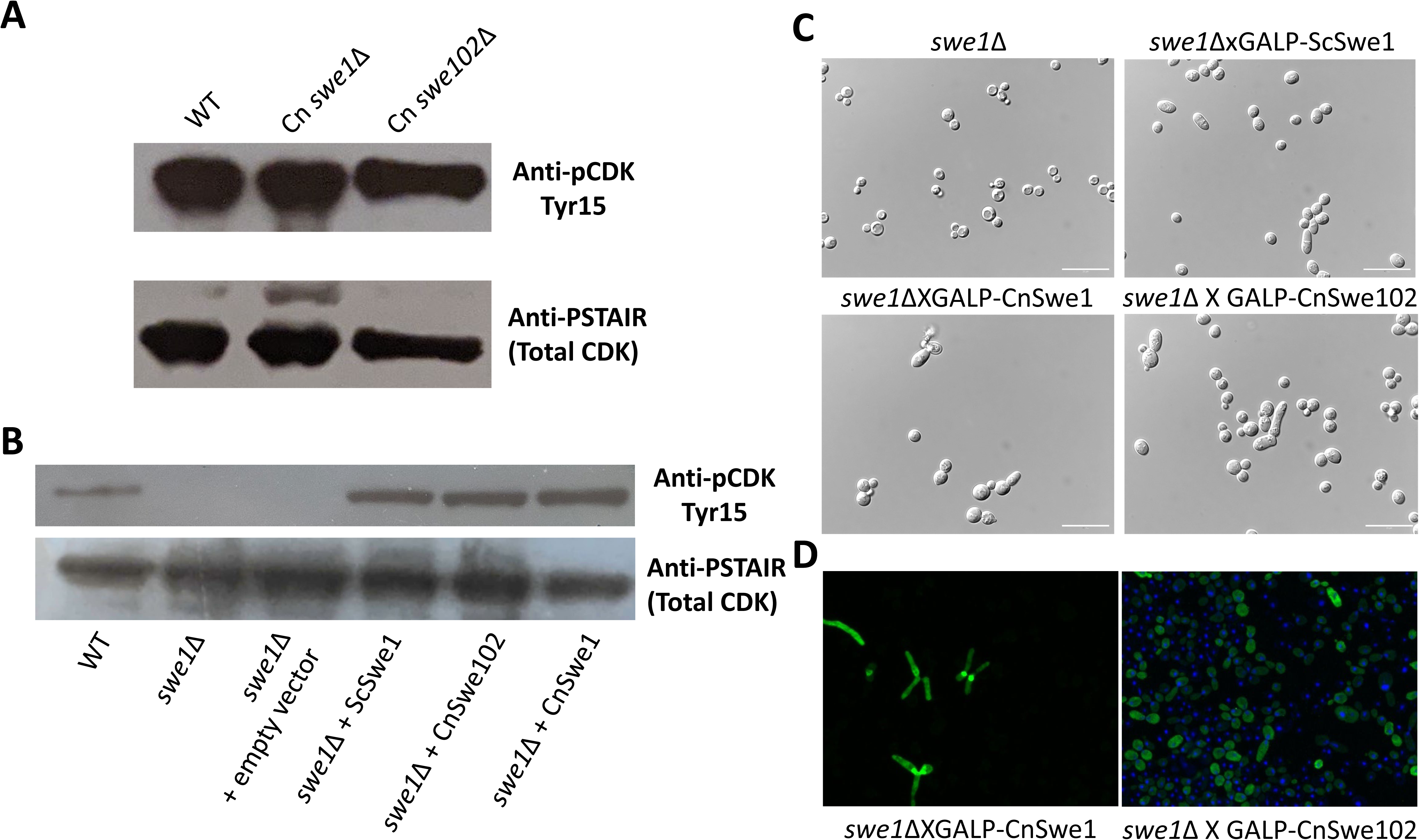
CnSwe1 and CnSwe102 promote CDK inhibitory phosphorylation and elongated morphology in *S. cerevisiae*. (A) Immunoblot analysis of CDK phosphorylation in WT, *swe1*Δ and *swe102*Δ strains of *C. neoformans*. CDK phosphorylation was detected using an anti-phospho-Cdc2 Tyr15 antibody, and anti-PSTAIR was used to detect total CDK. (B) CnSwe1 and CnSwe102 phosphorylate Cdc28 in *S. cerevisiae*. Phosphorylation of Cdc28 was detected using an anti-phospho-Cdc2 Tyr15 antibody in WT *S. cerevisiae*, *swe1*Δ cells carrying an empty vector, or *swe1*Δ cells expressing ScSwe1, CnSwe1 or CnSwe102 from plasmids. Anti-PSTAIR was used to detect total Cdc28. (C) Overexpression of ScSwe1, CnSwe1 or CnSwe102 induces elongated morphology in *S. cerevisiae*. *S. cerevisiae swe1*Δ cells carrying an empty vector or plasmids expressing the indicated Wee1-family kinases from the galactose-inducible GAL promoter were grown in galactose-containing medium and imaged by differential interference contrast (DIC) microscopy. (D) Cells expressing high levels of CnSwe1 or CnSwe102 exhibit elongated bud morphology. Cells treated as described in (C) were subjected to immunofluorescence microscopy to detect myc-tagged Wee1-family kinases using an anti-myc antibody. Scale bars, 20 μm.

To test whether CnSwe1 and CnSwe102 can function as Wee1-like CDK inhibitory kinases in a heterologous system, we expressed each protein in the *S. cerevisiae swe1*Δ background, which lacks the endogenous budding-yeast Wee1 homolog. Expression of either CnSwe1 or CnSwe102 restored Cdc28 tyrosine phosphorylation in *S. cerevisiae swe1*Δ cells, comparably to ScSwe1, as detected by anti-phosphotyrosine immunoblotting (Fig. 4B). Anti-PSTAIR detection of total Cdc28 served as a loading and CDK control. These results indicate that both cryptococcal Wee1-family kinases can phosphorylate the conserved CDK target in a heterologous budding-yeast system.

We next asked whether expression of the cryptococcal Wee1-family kinases produces the characteristic morphological consequence of increased Wee1 activity. Overexpression of ScSwe1 from the galactose-inducible GAL promoter induced elongated morphology in 63.6% of *S. cerevisiae swe1*Δ cells (n = 152), whereas fewer than 3% of vector-control *swe1*Δ cells were elongated. Under the same conditions, CnSwe102 and CnSwe1 induced elongated morphology in 47.7% (n = 131) and 64.4% (n = 118) of cells, respectively (Fig. 4C). Immunofluorescence microscopy using an anti-myc antibody confirmed that elongated cells contained high levels of myc-tagged CnSwe1 or CnSwe102 (Fig. 4D). Together, these results demonstrate that CnSwe1 and CnSwe102 retain conserved Wee1-like activity, as they promote Cdc28 phosphorylation and induce Wee1-associated elongated morphology when expressed in *S. cerevisiae*.

### *SWE1* and *SWE102* contribute to DNA-damaging stress tolerance in *C. neoformans*

Given that the *swe102*Δ P*_CTR4_*:*SWE1* strain was highly susceptible to HU and MMS in the extended stress screen and that CnSwe1 and CnSwe102 exhibited conserved CDK inhibitory activity associated with cell-cycle regulation, we reasoned that this phenotype may reflect a defect in replication- or DNA damage-associated checkpoint control (Elbaek et al., 2022; Koh, 2022). We hypothesized that these Wee1-family kinases play a role in the response to DNA damage and replication stress. We therefore directly examined the contribution of the CnSwe1–CnSwe102 module to DNA-damage and replication-stress tolerance. WT, P*_CTR4_*:*SWE1*, *swe102*Δ, and three independent *swe102*Δ P*_CTR4_*:*SWE1* strains were exposed to methyl methanesulfonate (MMS), camptothecin (CPT), or hydroxyurea (HU). Under conditions in which WT, P*_CTR4_*:*SWE1*, and *swe102*Δ strains retained growth, the *swe102*Δ P*_CTR4_*:*SWE1* strains showed marked sensitivity to MMS, CPT, and HU (Fig. 5A). This defect was reduced under BCS-treated conditions, in which expression of *SWE1* from the *CTR4* promoter is induced, suggesting that sufficient *SWE1* expression can partially compensate for the absence of *SWE102* during DNA-damaging stress.

**Fig. 5.**
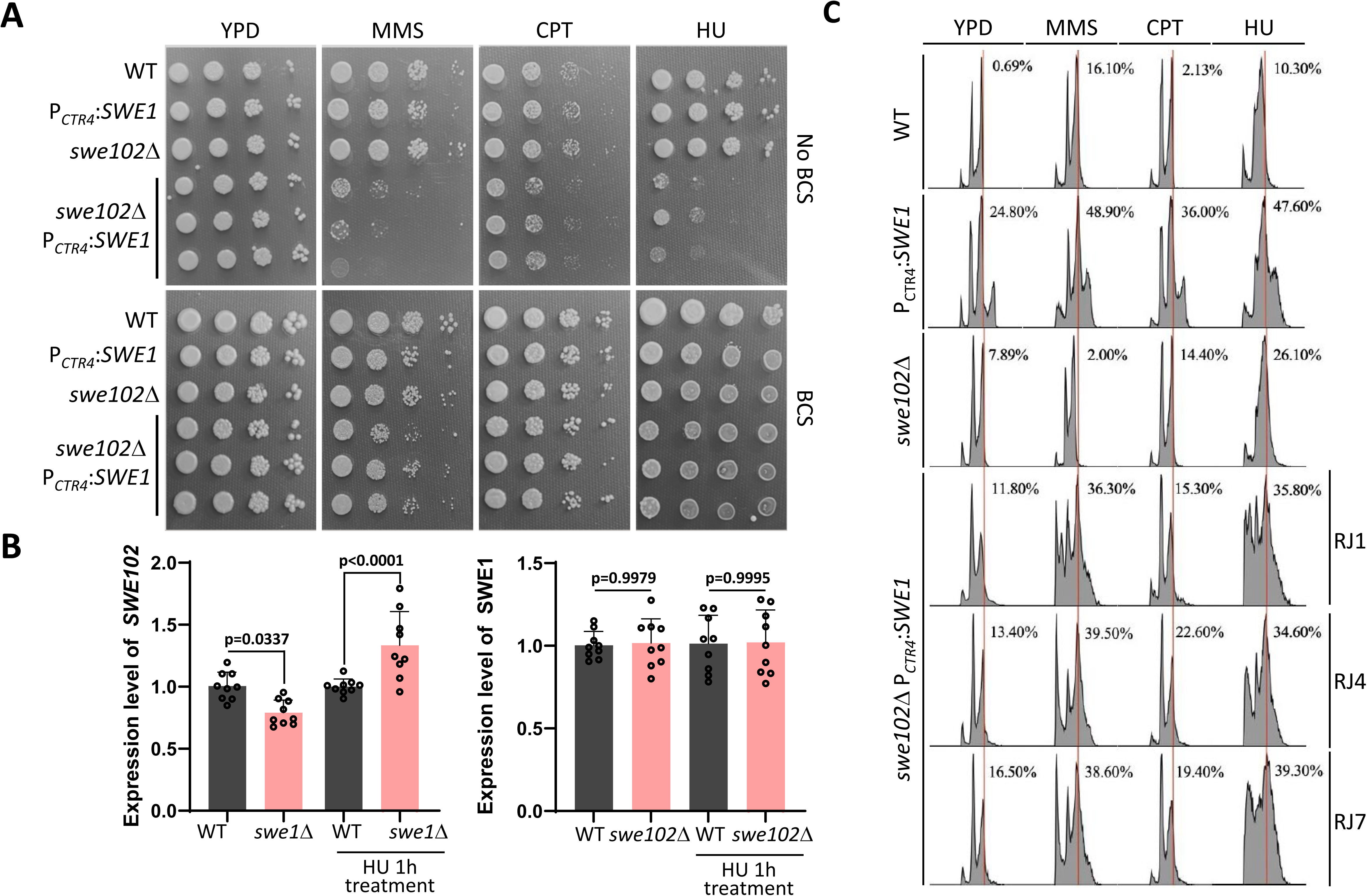
***SWE1* and *SWE102* contribute to tolerance against DNA-damaging stress in *C. neoformans*.** (A) Serial-dilution growth analysis of H99 WT (LK54), P*_CTR4_*:*SWE1*, *swe102*Δ (YSB1564), and three independent *swe102*Δ P*_CTR4_*:*SWE1* strains under DNA-damaging stress conditions. Cells were grown in liquid YPD medium, serially diluted 10-fold, and spotted onto YPD agar medium containing no drug, methyl methanesulfonate (MMS; 0.02%), camptothecin (CPT; 30 μM), or hydroxyurea (HU; 25 mM). (B) qRT–PCR analysis of reciprocal *SWE1* and *SWE102* expression in WT, *swe1*Δ, and *swe102*Δ strains under basal and HU-treated conditions. Overnight cultures were subcultured into fresh YPD medium at an initial OD_600_ of 0.2 and grown to OD_600_ 0.8. Cells were left untreated or treated with HU for 1 h before RNA extraction. *SWE1* transcript levels were analysed in WT and *swe102*Δ strains, and *SWE102* transcript levels were analysed in WT and *swe1*Δ strains. Transcript levels were normalized to *ACT1* and then to the corresponding WT control under each condition. Data are shown as mean ± SD. Statistical significance was determined by unpaired two-tailed Student’s t-test for each comparison. (C) DNA-content analysis of the strains shown in (A). Cells were cultured at 30 °C for 20 h in liquid YPD medium containing no drug, MMS, CPT, or HU at the concentrations described in (A). Cells were then stained with SYTOX Green and analysed by fluorescence flow cytometry. Percentage values indicate the fraction of the cell population with DNA content above 2n.

To determine whether reciprocal regulation between the two Wee1-family kinase genes contributes to this response, we measured *SWE1* and *SWE102* transcript levels under basal conditions and after short-term HU treatment. *SWE1* transcript levels were compared between WT and *swe102*Δ strains, whereas *SWE102* transcript levels were compared between WT and *swe1*Δ strains. Under basal conditions, loss of *SWE102* did not substantially alter *SWE1* expression, whereas *SWE102* expression was reduced in the *swe1*Δ mutant. After HU treatment, however, *SWE102* expression was significantly increased in the *swe1*Δ mutant compared with HU-treated WT cells (Fig. 5B). These results suggest that *SWE102* can be transcriptionally induced when *SWE1* is absent during replication stress.

We next assessed whether altered Wee1-family kinase dosage affected DNA-content profiles under DNA-damaging stress. Cells were cultured for 20 h in YPD medium containing MMS, CPT, or HU and then analyzed by flow cytometry following fixation and SYTOX Green staining. DNA-content analysis revealed an increased fraction of cells with DNA content above 2n in the P*_CTR4_*:*SWE1* and *swe102*Δ P*_CTR4_*:*SWE1* strains under DNA-damaging stress conditions (Fig. 5C). Together, these results indicate that the functional balance between *SWE1* and *SWE102* is particularly important under genotoxic and replication-stress conditions, supporting a role for cryptococcal Wee1-family kinases in maintaining proper DNA replication and cell-cycle control.

### *SWE1* and *SWE102* contribute to cryptococcal virulence in a murine infection model

We next assessed whether the stress-adaptation and cell-cycle-related functions of CnSwe1 and CnSwe102 contribute to cryptococcal virulence *in vivo*. In a murine intrapharyngeal infection model, mice infected with the *swe1*Δ mutant were essentially avirulent compared with those infected with the wild-type strain (Fig. 6A). Whereas WT-infected mice succumbed rapidly to infection, *swe1*Δ-infected mice survived throughout the observation period, indicating that loss of *SWE1* nearly abolished cryptococcal virulence under these conditions. This defect was largely restored by reintroduction of *SWE1*-mCherry, confirming that the virulence attenuation was attributable to loss of *SWE1*. The *swe102*Δ mutant also showed significantly attenuated virulence relative to WT, although the effect was less severe than that observed for *swe1*Δ. Complementation with *SWE102*-mCherry restored survival kinetics toward the WT pattern (Fig. 6A).

**Fig. 6.**
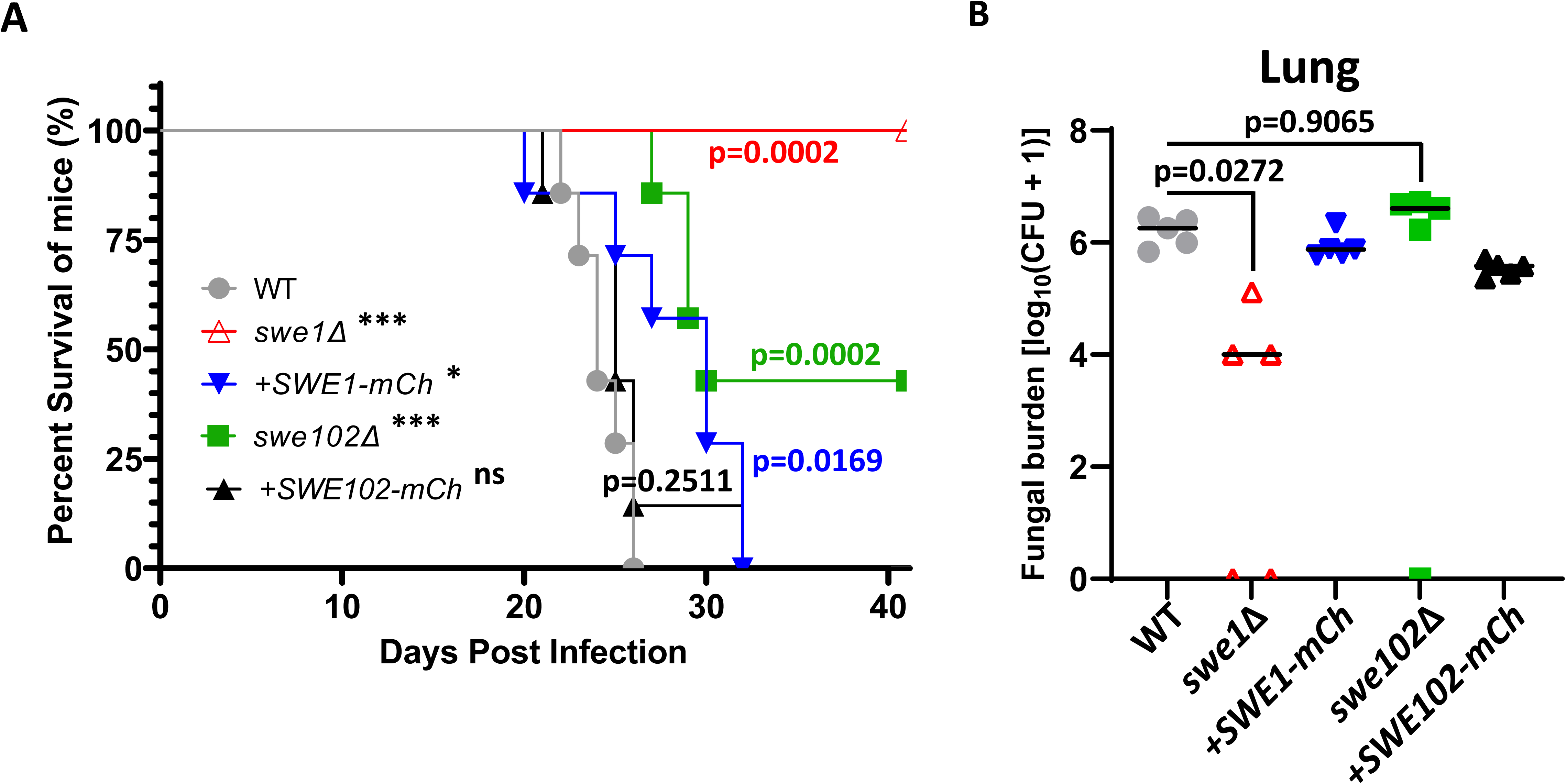
***SWE1* and *SWE102* contribute to cryptococcal virulence in a murine infection model.** (A) Survival of mice infected with WT, *swe1*Δ, *swe1*Δ::*SWE1*-mCherry, *swe102*Δ, or *swe102*Δ::*SWE102*-mCherry strains. Seven-week-old male C57BL/6J mice were infected intrapharyngeally with 5 × 10 cells in 20 μl PBS and monitored for survival. Survival data are shown as Kaplan–Meier curves. Statistical significance was determined by the log-rank Mantel–Cox test. (B) Pulmonary fungal burden in mice infected with WT, *swe1*Δ, *swe1*Δ::*SWE1*-mCherry, *swe102*Δ, or *swe102*Δ::*SWE102*-mCherry strains. Lungs were harvested at 21 days post-infection, homogenized, serially diluted, and plated on YPD medium to determine colony-forming units (CFU). Statistical significance was determined as indicated in the panel.

Consistent with the survival data, pulmonary fungal burden analysis at 21 days post-infection revealed a marked reduction in lung colonization by the *swe1*Δ mutant (Fig. 6B). WT-infected mice carried high fungal burdens in the lung, whereas *swe1*Δ-infected mice showed substantially reduced fungal loads, with some mice yielding no detectable CFU. Reintroduction of *SWE1*-mCherry restored lung fungal burden toward WT levels. By contrast, the *swe102*Δ mutant did not show a significant reduction in pulmonary fungal burden at this time point, despite its attenuated survival phenotype. Together, these results indicate that both *SWE1* and *SWE102* contribute to cryptococcal virulence *in vivo*, with *SWE1* playing a particularly important role in pulmonary colonization.

## DISCUSSION

In this study, we found that the human fungal pathogen *C. neoformans* encodes two divergent Wee1-family kinases, CnSwe1 and CnSwe102, that retain conserved CDK-inhibitory activity but function outside the canonical *S. cerevisiae* morphogenesis checkpoint. Neither kinase concentrated at the mother-bud neck, and the two acted in an interdependent, dosage-sensitive manner: altering *SWE1* dosage in the absence of *SWE102* sensitized cells to replication- and DNA-damaging stresses, and the double mutant could not be recovered. Importantly, loss of *SWE1* nearly abolished virulence in a murine infection model. Together, these findings indicate that the *C. neoformans* Wee1–CDK module acts as a dosage-sensitive checkpoint that promotes replication-stress adaptation and fungal virulence, which is distinct from septin-dependent morphogenesis checkpoint as described in *S. cerevisiae*.

Our phylogenetic analysis points to three scenarios with respect to the number of WEE1-encoding genes. Model fungal ascomycete species possess a single WEE1-encoding gene. Vertebrates, including humans, encode two Wee1 kinases that are closely related yet functionally specialized with one operating in somatic cells and the other devoted to germline biology (Han and Conti, 2006). Our analysis reveals a third distinct case that occurs in basidiomycete yeasts of the order Tremellales, including *C. neoformans*. These species encode two phylogenetically divergent forms of Wee1 kinase. It remains unknown how prevalent this third scenario is within the tree of life. Interestingly, while *Trypanosoma brucei* encodes a single WEE1 homolog, *T. cruzi* genome contains two highly divergent WEE1 homologs (Boynak et al., 2013), although their specific roles and the significance of possessing two divergent WEE1-encoding genes in this species have not been investigated. The presence of two distinct WEE1 proteins in *C. neoformans* does not appear to be specific to pathogenicity, given that some non-pathogenic Tremellales species also contain both WEE1-encoding genes in their genomes.

Based on the investigation presented here, the two WEE1 homologs in *C. neoformans* operate during vegetative growth yet their functions do not completely overlap. Instead, they likely play complementary and partly overlapping roles in cell cycle regulation under various stress conditions. The difficulty in obtaining the *swe1*Δ *swe102*Δ double mutant suggests that the two WEE1 homologs are collectively essential. However, the *swe102*Δ P*_CTR4_*:*SWE1* strain proliferates under conditions that markedly reduce *SWE1* expression in the absence of *SWE102*, albeit with a slight growth defect (Fig. 3). A likely explanation for this apparent contradiction is that *CTR4*-mediated repression is incomplete: CuSO_4_ strongly reduces but does not fully eliminate *SWE1* transcription (Fig. 3B), so the residual CnSwe1 produced under repressive conditions may be sufficient to support viability in the absence of *SWE102*. A non-exclusive possibility is that the two homologs are collectively essential only under stress conditions, such that the inability to recover the double mutant reflects the stress associated with transformation rather than an absolute growth requirement under standard conditions. Distinguishing between these scenarios will require a fully inactivated or conditionally degraded allele, for example, a complete deletion combined with an inducible-degron system to test whether near-complete loss of CnSwe1 activity is lethal in the *swe102*Δ background.

Results of the heterologous expression of CnSwe1 and CnSwe102 in *S. cerevisiae* are consistent with both kinases acting as Cdk1 inhibitors in *C. neoformans* under various circumstances. Future studies will define their precise contributions to specific cell-cycle checkpoints, although the data presented here suggest that both proteins act in DNA-damage and replication checkpoints. Interestingly, the *S. cerevisiae* mutants with defects in Swe1 degradation show HU sensitivity because of high Swe1 protein levels and not because of DNA damage accumulation or incomplete DNA synthesis; instead, the sensitivity of those mutants result from dramatically delayed recovery from HU-induced S-phase arrest (Liu and Wang, 2006). Moreover, an *S. cerevisiae* mutant lacking Swe1 is not hypersensitive to HU (Matmati et al., 2009). Thus, unlike in *S. cerevisiae*, the two WEE1 homologs in *C. neoformans* appear to play a role similar to that in mammalian cells, where they regulate replication during genotoxic stress (Vakili-Samiani et al., 2022).

In *S. cerevisiae*, timely Swe1 degradation that allows progression to mitosis requires its recruitment to the septin complex assembled at the mother-bud neck, which in turn depends on the Hsl1 and Elm1 kinases and on recruitment of the Hsl7 protein to the septin ring (Kang et al., 2016; Szkotnicki et al., 2008). Phosphorylation of ScSwe1 by Cdc28 and the polo kinase Cdc5 occurs at multiple sites, which triggers the degradation required for cell-cycle progression. Disruption of the actin cytoskeleton or perturbations in septin organization at the mother-bud neck trigger Swe1-dependent inhibition of mitosis known as the morphogenesis checkpoint (Keaton and Lew, 2006; Lee et al., 2005). Strikingly, *C. neoformans* cells lacking the septin complex at the mother-bud neck proliferate under non-stress conditions (Kozubowski et al., 2009). This suggests that *C. neoformans* either lacks morphogenesis checkpoint or possess one that does not depend on the septin complex at the mother-bud neck. Thus, while our data do not exclude a morphogenesis checkpoint that senses bud formation in *C. neoformans*, our results suggest that Swe1 and Swe102 are not regulated through septin-complex formation, in contrast to ascomycetous yeasts.

Data presented here, together with previously published work, suggest that both Swe1 and Swe102 contribute to the pathogenicity of *C. neoformans*, with Swe1 playing a critical role during infection—consistent with these kinases being important for survival at various stress conditions *in vitro*. Given that WEE1 is an attractive anticancer target, developing or repurposing anticancer drugs that inhibit *C. neoformans* Swe1/Swe102 may represent a promising anticryptococcal strategy. Future studies will explore the mechanistic details of how these WEE1 homologs contribute to stress survival and investigate how these mechanisms can be leveraged to better treat cryptococcosis.

## MATERIALS AND METHODS

### Ethics statement

All animal experiments were performed in accordance with institutional and national guidelines for animal care and use. Animal protocols were reviewed and approved by the Institutional Animal Care and Use Committee of Hallym University under approval number HallymR1 (2024-6). Mice were monitored at least twice daily after infection, and humane endpoints were applied according to the approved protocol.

### Strains and growth conditions

*S. cerevisiae* strains were routinely cultured in yeast extract–peptone–dextrose (YPD) medium or yeast extract–peptone–adenine–dextrose (YPAD) medium at 30 °C. YPD medium contained 1% yeast extract, 2% Bacto peptone and 2% glucose. YPAD medium was prepared by supplementing YPD with adenine hemisulfate at 100 mg/l. Solid media were prepared by adding 2% Bacto agar. For lithium acetate/single-stranded carrier DNA/polyethylene glycol transformation, double-strength 2× YPAD medium was used where indicated, and solid media were prepared with 2% Bacto agar. *C. neoformans* strains were routinely cultured in liquid or solid YPD medium at 30 °C with shaking at 220 rpm unless otherwise indicated. Overnight cultures were typically grown for 16–18 h and used for strain construction, phenotypic assays, protein extraction, microscopy, flow cytometry and animal infection experiments.

### Bioinformatic, phylogenetic and structural analyses of Wee1-family kinases

Predicted Wee1-family kinase homologs were identified from representative fungal and eukaryotic species using annotated protein sequences retrieved from public genome databases, including FungiDB, NCBI, UniProt and organism-specific genome resources where applicable. The *C. neoformans* H99 Wee1-family proteins CnSwe1 and CnSwe102 were identified as CNAG_03171 and CNAG_03369, respectively. Representative Wee1-family proteins from *C. neoformans* H99, *C. deneoformans* JEC21, *C. gattii* R265, *Kwoniella heveanensis*, *Tremella mesenterica*, *Ustilago maydis*, *Saccharomyces cerevisiae*, *Schizosaccharomyces pombe*, *Drosophila melanogaster*, and *Homo sapiens* were used for comparative analysis.

Full-length amino acid sequences were aligned using ClustalX2, and the phylogenetic tree was generated and visualized using the online tool iTOL (Interactive Tree of Life; https://itol.embl.de/). The resulting tree was used to determine the evolutionary relationship of CnSwe1 and CnSwe102 to representative fungal and eukaryotic Wee1-family kinases. Protein domain architecture was analysed using InterPro. Protein lengths and predicted Wee1-like kinase domains were visualized for representative fungal Wee1-family proteins. Conserved kinase-associated sequence motifs in CnSwe1 and CnSwe102 were aligned using ClustalX2. The aligned regions included the ATP-binding/glycine-rich region, Lys-containing kinase subdomain, catalytic HxD-like region, activation-segment motif, and APE/CPE-like motif. Residue positions were assigned according to the full-length CnSwe1 and CnSwe102 amino acid sequences. AlphaFold3-predicted structures of CnSwe1 and CnSwe102 were generated using AlphaFold3 through the AlphaFold Server (https://alphafoldserver.com/). For structural comparison, the C-terminal kinase-core regions corresponding to CnSwe1 residues 795–1084 and CnSwe102 residues 818–1146 were extracted and superimposed using UCSF ChimeraX MatchMaker. Structural similarity was assessed by calculating the root-mean-square deviation (RMSD) over pruned Cα atom pairs. Per-residue pLDDT confidence values were extracted from the AlphaFold prediction files and plotted across the full-length proteins. Mean pLDDT values were calculated for both full-length proteins and the defined kinase-core regions.

### Construction of *C. neoformans* deleted mutant, complemented and promoter-replacement strains

To generate *swe1*Δ mutants, the *SWE1* locus was disrupted in the *C. neoformans* serotype A *MAT*α wild-type strain H99 by homologous recombination using a gene disruption cassette containing the nourseothricin acetyltransferase resistance marker (*NAT*). Mutant construction was performed according to previously described *C. neoformans* genetic manipulation methods (Kim et al., 2009). Briefly, the gene disruption cassette was introduced into H99 by biolistic transformation, and transformants were selected on YPD agar supplemented with nourseothricin (100 mg/L). Candidate *swe1*Δ mutants were screened by diagnostic PCR, and correct targeted gene deletion was confirmed by Southern blot analysis, as shown in Supplementary Fig. 2.

To validate the observed phenotypes of the *swe1*Δ and *swe102*Δ mutants and to examine the subcellular localization of Swe1 and Swe102, mCherry-tagged complemented strains were constructed using Gibson assembly. Full-length *SWE1* and *SWE102* gene fragments, including their native regulatory regions, were amplified from H99 genomic DNA by Phusion PCR. The amplified fragments were cloned into the pNEO-mCherry plasmid. Correct insertion of each target gene into the plasmid was confirmed by restriction enzyme digestion and sequencing analysis. For *SWE1* complementation, the plasmid carrying *SWE1-mCherry* was linearized with AfeI and introduced into the *swe1*Δ mutant strain by biolistic transformation. For *SWE102* complementation, the plasmid carrying *SWE102-mCherry* was linearized with PacI and introduced into the *swe102*Δ mutant strain by biolistic transformation. Stable transformants were selected on appropriate drug-containing medium, and correct targeted reintegration into the native locus was confirmed by diagnostic PCR. Functionality of the mCherry-tagged alleles was assessed by complementation of mutant growth phenotypes under selected stress conditions.

To generate strains in which *SWE1* expression is regulated by the copper-responsive *CTR4* promoter, P*_CTR4_*:*SWE1* promoter-replacement strains were constructed following established *CTR4* promoter-replacement protocols (Ory et al., 2004). The left flanking region upstream of the SWE1 start codon and the 5′ region of the SWE1 open reading frame were amplified from H99 genomic DNA by PCR. The NAT-P*_CTR4_*cassette was amplified from the pNAT-*CTR4*-2 plasmid. These fragments were assembled by overlap PCR and introduced into *C. neoformans* by biolistic transformation. Transformants were selected on YPD agar supplemented with nourseothricin (100 mg/L), and correct insertion of the P*_CTR4_*:*SWE1* cassette was verified by diagnostic PCR. P*_CTR4_*:*SWE1* strains were generated in both wild-type and *swe102*Δ genetic backgrounds. For promoter regulation experiments, P*_CTR4_:SWE1* expression was repressed with 25 μM CuSO_4_ and induced with the copper chelator bathocuproinedisulfonic acid (BCS; 200–300 μM).

### Generation of *Saccharomyces cerevisiae* strains expressing *C. neoformans* WEE1 homologues

*S. cerevisiae* strains expressing *C. neoformans* homologues of WEE1 from the GAL4 promoter were based on *swe1*Δ mutant strain DLY1028 (Sia et al., 1996) that was transformed with expression constructs based on plasmid pJM1101, which originally expressed *S. cerevisiae SWE1* allele (McMillan et al., 2002). Plasmid pJM1101 was modified by swapping *S. cerevisiae SWE1* sequence with *CnSWE1* or *CnSWE102* intron-less coding sequences which were codon-optimized for expression in *S. cerevisiae*. Swapping sequences was performed by homologues recombination by GenScript. Similarly, *S. cerevisiae* strain DLY1028 was transform with constructs expressing the two *C. neoformans* homologues of WEE1 driven by the endogenous *S. cerevisiae SWE1* promoter based on plasmid pJM1115 (McMillan et al., 2002).

### Growth and stress susceptibility assays

*C. neoformans* strains were routinely cultured in YPD medium at 30 °C for 16–18 h with shaking at 220 rpm. For serial-dilution growth assays, overnight cultures were diluted into fresh YPD medium and grown to approximately 1 × 10 cells/ml. Cells were washed where necessary, adjusted to equivalent densities, serially diluted 10-fold, and spotted onto solid medium using 3–5 μl of each dilution. Plates were incubated under the indicated conditions and photographed after 1–5 days depending on the growth rate and stress condition.

For selected stress profiling of *SWE1* and *SWE102* mutant strains, cells were spotted onto YPD agar and incubated under temperature and CO_2_ stress conditions or onto YPD agar supplemented with stress-inducing agents. Conditions included growth at 30°C, 37°C, 37°C with 5% CO_2_, and 39°C. Chemical stress conditions included diamide (2 mM), tunicamycin (0.3 μg/ml), NaCl (1.5 M), fludioxonil (1 μg/ml), fluconazole (10 μg/ml), sodium dodecyl sulfate (0.03%), and other stressors as indicated in the corresponding figure legends. For extended stress profiling, plates were supplemented with the indicated cell wall, membrane, osmotic, antifungal, oxidative, heavy metal, ER, and genotoxic stress-inducing agents. Plates were photographed after 3–4 days of incubation unless otherwise indicated.

For copper-dependent regulation assays, WT, P*_CTR4_*:*SWE1*, and *swe102*Δ P*_CTR4_*:*SWE1* strains were spotted onto YPD agar, YPD agar supplemented with CuSO_4_, or YPD agar supplemented with BCS. For dose-dependent regulation assays, plates contained CuSO_4_ at 25, 50, or 100 μM, or BCS at 200, 300, or 600 μM, as indicated. Plates were incubated at 30°C and photographed after 3–4 days.

For pH-dependent growth assays, overnight cultures were grown in YPD medium at 30°C, serially diluted 10-fold, and spotted onto unbuffered RPMI agar or RPMI agar adjusted to pH 5, pH 7, or pH 8. The pH of RPMI medium was adjusted before sterilization using 50 mM citrate (citric acid/NaOH) for pH 5 and 150 mM HEPES for pH 7 or pH 8, with final adjustment using NaOH. Plates were incubated at 37°C in ambient air or at 37°C with 5% CO and photographed after the indicated incubation period. These assays were used to compare growth of WT, *swe1*Δ, *swe102*Δ, *swe1*Δ::*SWE1*-mCherry, and *swe102*Δ::*SWE102*-mCherry strains under pH-adjusted and CO -dependent conditions.

### Fluorescence microscopy of *C. neoformans* mCherry-tagged strains

For localization analysis of CnSwe1-mCherry and CnSwe102-mCherry, *swe1*Δ::*SWE1*-*mCherry* and *swe102*Δ::*SWE102*-*mCherry* strains were grown overnight in YPD medium at 30 °C. Overnight cultures were subcultured into fresh YPD medium at an initial OD of 0.2 and grown at 30 °C to OD 0.8. Cells were left untreated or treated with fludioxonil (1 μg/ml) or fluconazole (10 μg/ml) for 1 h. After treatment, cells were collected by centrifugation, washed with PBS or sterile water, and stained with Hoechst to visualize nuclei. Cells were mounted on microscope slides and imaged by differential interference contrast and fluorescence microscopy using a Zeiss Axio Imager A2 microscope equipped with an AxioCam 506 color digital imaging system. DIC, mCherry, Hoechst, and merged images were acquired using identical or comparable exposure settings within each experiment and analysed using Fiji/ImageJ.

### RNA extraction and qRT–PCR analysis

For qRT–PCR analysis, cells were grown under the indicated conditions and harvested for total RNA extraction. For P*_CTR4_*:*SWE1* regulation assays, *swe102*Δ P*_CTR4_*:*SWE1* cells were grown in YPD medium, YPD medium supplemented with CuSO_4_, or YPD medium supplemented with BCS. For basal *SWE1* expression analysis, WT and *swe102*Δ P*_CTR4_*:*SWE1* strains were grown in YPD medium. For reciprocal *SWE1* and *SWE102* expression analysis under replication stress, overnight cultures were subcultured into fresh YPD medium at an initial OD of 0.2 and grown to OD 0.8. Cells were then left untreated or treated with HU for 1 h before RNA extraction. Cells were harvested by centrifugation, rapidly frozen in liquid nitrogen, and lyophilized where applicable. Total RNA was extracted using the Easy-BLUE RNA extraction kit according to the manufacturer’s instructions, with minor modifications as described previously. RNA samples were treated with DNase where applicable to remove contaminating genomic DNA. cDNA was synthesized from 1 μg total RNA using reverse transcriptase. qRT–PCR was performed using gene-specific primers for *SWE1*, *SWE102*, and *ACT1*. Transcript levels were normalized to *ACT1* and quantified using the comparative Ct method. For each experiment, transcript levels were normalized to the corresponding WT or untreated control as indicated in the figure legend. Data represent the mean ± SD from biological replicates.

### Morphology analysis and cell-length quantification

For morphology analysis, the indicated *C. neoformans* strains were grown on YPD or DME medium with or without CuSO_4_ or BCS, as indicated. Cells were collected from colonies after the indicated incubation period, resuspended in sterile water or PBS, and examined by differential interference contrast microscopy using a Zeiss Axio Imager A2 microscope equipped with an AxioCam 506 color digital imaging system. Images were analysed using Fiji/ImageJ. For round yeast cells, cell diameter was measured. For elongated cells or incompletely separated cells, the maximum cell length was measured. Cell-length measurements were obtained from 50 cells per strain and condition across three independent biological replicates. Statistical significance was determined using GraphPad Prism. For morphology quantification, ordinary two-way ANOVA followed by Tukey’s multiple-comparisons test was used unless otherwise indicated.

### Virulence factor-associated phenotype assays

To assess capsule-associated phenotypes, the indicated strains were spotted onto DME agar medium with or without CuSO_4_ or BCS and incubated at the indicated temperature for 3 days. Cells were collected from colonies, resuspended in sterile water, and examined microscopically where applicable. To assess melanin production, strains were spotted onto Niger seed agar medium with or without CuSO_4_ or BCS and incubated for 3 days. Plates were photographed using identical imaging settings within each experiment.

### Galactose induction in Saccharomyces cerevisiae

*S. cerevisiae* trains were grown overnight in YPAD medium at 30 °C with shaking at 225 rpm. After overnight incubation, cultures were diluted 1:5 into yeast extract–peptone–adenine medium supplemented with 2% raffinose (YPAR). This condition does not induce or repress the GAL4 promoter but permits binding of regulatory factors required for subsequent galactose induction. When cultures reached late logarithmic phase, galactose was added to a final concentration of 2%. Cultures were then incubated for at least 6 h before cells were collected for immunofluorescence microscopy and protein extraction.

### Protein extraction and immunoblotting

Protein extraction was performed using a trichloroacetic acid (TCA)-based protocol. Log-phase cells were harvested by centrifugation for 3 min at approximately 10,000 × g. The supernatant was discarded, and the cell pellet was resuspended in 225 μl pronase buffer containing 1.4 M sorbitol, 25 mM Tris-HCl pH 7.5, 20 mM NaN and 2 mM MgCl . To each sample, 56 μl of 85% TCA solution was added, and the mixture was transferred to cryotubes containing 280 μl acid-washed glass beads (425–600 μm). Cells were disrupted by bead beating for eight cycles of 20 s each, with 2-min cooling intervals at 4 °C between cycles. After lysis, samples were chilled on ice for 10 min and transferred to new tubes. The beads were washed twice with 5% TCA, and the washes were combined with the lysates. Samples were incubated on ice for 5 min and centrifuged at 11,000 rpm for 10 min at 4 °C. The supernatant was discarded, and the protein pellets were resuspended in 30 μl Thorner buffer containing 8 M urea, 5% SDS, 40 mM Tris-HCl pH 6.8, 0.1 mM EDTA and 0.4 mg/ml bromophenol blue. When samples turned yellow or green, up to 10 μl of 2 M Tris base was added until the solution turned blue. β-mercaptoethanol was then added to a final concentration of 1% to denature proteins.

Samples were loaded onto NuPAGE 4–12% Bis-Tris gels (Invitrogen, NP0335BOX) and electrophoresed at 150 V for 55 min using MOPS running buffer containing 0.1 M MOPS, 0.1 M Tris base, 6.93 mM SDS and 2.05 mM EDTA. Proteins were transferred to PVDF membranes (Immobilon Transfer Membranes, Merck Millipore, IPFL07810) using 1× NuPAGE transfer buffer (Thermo Fisher Scientific, NP0006). Protein transfer was confirmed by Ponceau S staining. Membranes were washed twice with TBST for 30 min, blocked with 5% BSA in TBST for 1 h at room temperature and incubated overnight at 4 °C with primary antibodies. Anti-phospho-Cdc2 Tyr15 antibody (Cell Signaling Technology, 9111) was used to detect inhibitory CDK phosphorylation, and anti-PSTAIR antibody (Sigma, P7962) was used to detect total Cdk1 protein. HRP-conjugated secondary antibodies were used, and signals were developed using SuperSignal West Pico Chemiluminescent Substrate (Thermo Fisher Scientific, 34580).

### DNA-content analysis by flow cytometry

For DNA-content analysis, approximately 1 × 10^7^ cells were collected by centrifugation at 13,000 × g, and the supernatant was discarded. Cells were fixed overnight in 70% ethanol at 4 °C. The following day, fixed cells were washed with 1× citrate solution containing 0.890 μM citric acid and 0.491 M sodium citrate. Cells were then centrifuged and resuspended in 1 ml 1× citrate solution supplemented with RNase A (0.25 mg/ml) and SYTOX Green (500 nM). Samples were incubated overnight at 4 °C in the dark and sonicated briefly before flow cytometric analysis. DNA content was analyzed using a CytoFLEX flow cytometer (Beckman Coulter Life Sciences).

### Immunofluorescence microscopy of *S. cerevisiae* cells

For immunofluorescence microscopy, *S. cerevisiae* strains were cultured overnight in 5 ml YPAD medium at 30 °C with shaking at 220 rpm. Cultures were diluted 1:20 into fresh YPAD medium and grown at 30 °C until they reached approximately 1–2 × 10 cells/ml. At this density, 0.5 ml of 37% formaldehyde was added directly to each culture, and cells were fixed for 2 h at 30 °C with shaking at 200 rpm. Fixed cells were harvested by centrifugation for 2 min at 1,500 × g, washed with 1× PBS and centrifuged again. Cell pellets were washed with solution A containing 100 mM KPO pH 6.5, 0.5 mM MgCl and 1.2 M sorbitol. Cells were treated with 10 μl β-mercaptoethanol and 50 μl lyticase solution (10 mg/ml; Zymolyase 100T concentrate in 0.1 M sorbitol; US Biological, Z100) and incubated at 37 °C for 20 min. Spheroplasted cells were then washed twice with solution A and once with 1× PBS, followed by resuspension in 600 μl 1× PBS containing 1% BSA. Immunofluorescence slides (Polysciences, 183571) were prepared by applying 5 μl of 0.1% poly-L-lysine (Sigma, P1524) to each well. After 30 s, poly-L-lysine was aspirated, and each well was washed three times with distilled water. Five microliters of lyticase-treated cells were added to each well and allowed to attach for 30–60 s. Cells were aspirated, and slides were air-dried until the wells became opaque, approximately 5 min. Cells were permeabilized with 2 μl of 0.1% SDS for 5 min at room temperature and washed eight times with 1× PBS containing 1% BSA. Cells were then incubated with 5 μl primary antibody solution in a humidified chamber overnight at 4 °C. After eight washes with 1× PBS containing 1% BSA, 5 μl secondary antibody solution was added, and slides were incubated for 1 h at room temperature in the dark. Antibody solution was aspirated, and wells were washed eight times with 1× PBS containing 1% BSA. Before the wells dried, 5 μl ProLong Glass Antifade Mountant with NucBlue stain (Thermo Fisher Scientific, P36981) was applied to each well, and slides were coverslipped. Slides were pressed for 5 min, incubated for 14–16 h at room temperature in the dark and then imaged. Microscopic images were acquired using a Leica DMi8 inverted microscope (Leica Microsystems) and Leica Application Suite X software. Images were analyzed using Fiji/ImageJ. Cell counts and morphology measurements were performed using Fiji.

### In vivo virulence assay

The wild-type strain (H99), the *swe1*Δ mutant (YSB8309), the *swe1*Δ+*SWE1* complemented strain (YSB9411), the *swe102*Δ mutant (YSB1564), and the *swe102*Δ+*SWE102* complemented strain (YSB8121) were grown overnight in YPD broth. The yeast cells obtained were then pelleted and resuspended in sterile PBS to achieve a concentration of 2.5 × 10^6^ cells/ml, as determined by hemocytometer counting. Groups of 15 male C57BL/6J mice, aged 7 weeks (Dooyeol Biotech), were anesthetized using isoflurane. The mice were infected intrapharyngeally with 5 × 10^4^ cells in 20 μl of PBS. The cell concentration in the inoculum was verified by plating serial dilutions and counting the CFU. The mice were observed twice daily. For the fungal burden assay, five mice from each group were euthanized by CO_2_ inhalation 21 days after infection. The lungs and brain were collected and homogenized in 1 ml of PBS, and serial dilutions were plated on YPD medium to measure the CFU. Additionally, three mice from each group were euthanized by CO_2_ inhalation 21 days post-infection, and their lungs and brains were collected for histopathological analysis. The log-rank (Mantel-Cox) test was used to assess differences in survival curves, with P-values <0.05 considered significant.

## Acknowledgements

We thank Dr. Daniel Lew (MIT) for providing the *S. cerevisiae swe1*Δ strain DLY1028 that served as a heterologous host to express CnSwe1 and CnSwe102 allelles, and the plasmids pJM1101 and pJM1115.

## Data availability

The original contributions presented in the study are included in the article and Supplementary Material. Additional raw data supporting the conclusions of this article, including source data for growth assays, microscopy, qRT–PCR, immunoblotting, flow cytometry, and animal infection experiments, will be made available by the corresponding authors upon reasonable request, without undue reservation.

## Author contributions

J-TC, RJC-R, LK, and Y-SB conceived and designed the study. J-TC, RJC-R, D-HY, S-HL, SE, and NB performed experiments and/or analyzed data. J-TC, RJC-R, KTS, SC, LK, and Y-SB interpreted the results. J-TC and RJC-R drafted the manuscript with input from LK and Y-SB. LK and Y-SB supervised the study. All authors contributed to manuscript revision and approved the submitted version.

## Supplementary material

The Supplementary Material for this article includes Supplementary Figures 1–5 and Supplementary Tables 1–2.

---

**Supplementary Fig. 1.**
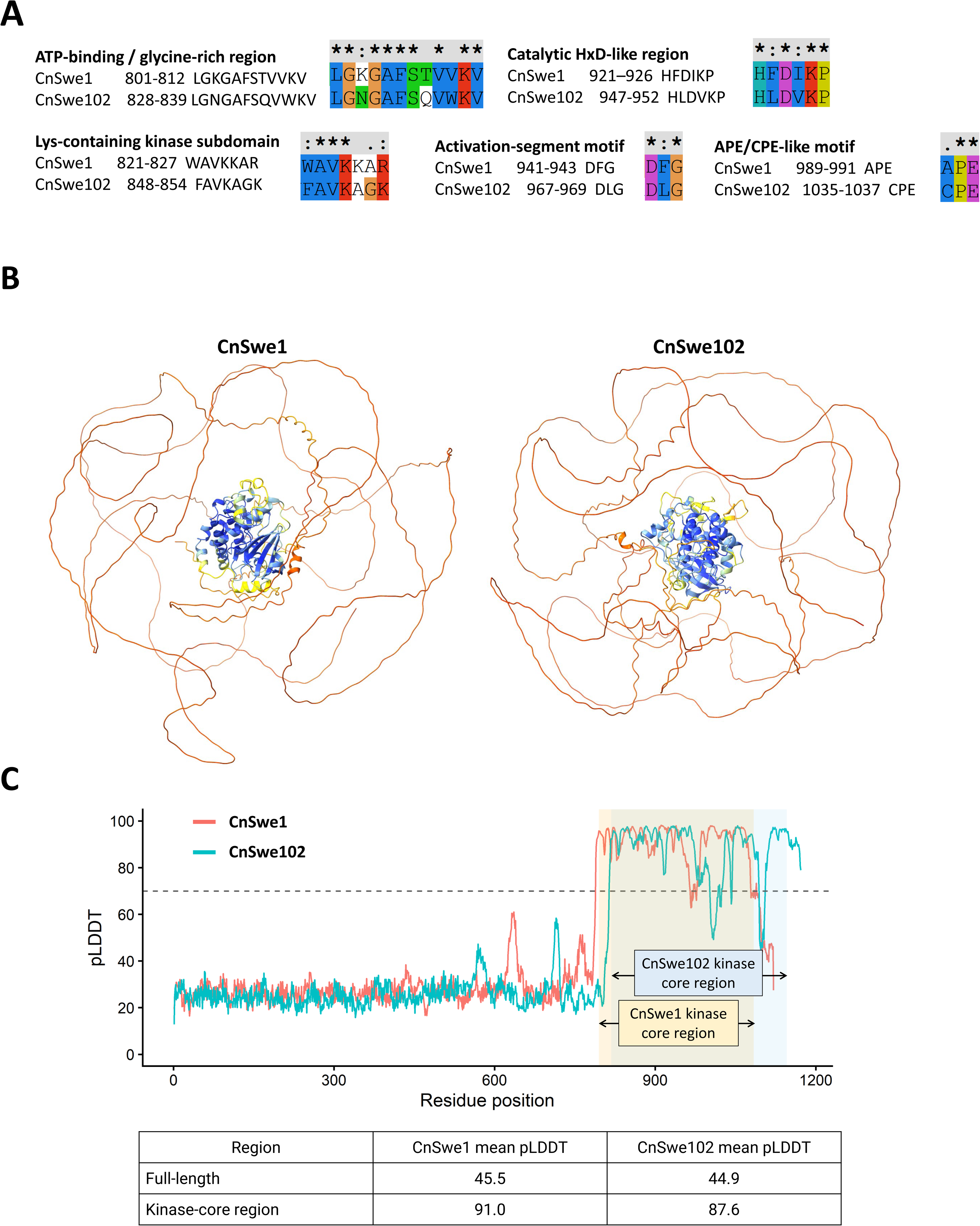
Sequence motif conservation and AlphaFold prediction-confidence analysis of CnSwe1 and CnSwe102. (A) Alignment of representative conserved kinase-associated motifs in CnSwe1 and CnSwe102. The indicated regions include the ATP-binding/glycine-rich region, Lys-containing kinase subdomain, catalytic HxD-like region, activation-segment motif, and APE/CPE-like motif. Residue positions correspond to full-length protein coordinates. Sequence alignment was performed using ClustalX2. (B) Full-length AlphaFold-predicted structures of CnSwe1 and CnSwe102 shown with pLDDT confidence coloring. (C) AlphaFold-derived per-residue pLDDT profiles of full-length CnSwe1 and CnSwe102. The shaded regions indicate the kinase-core regions used for structural comparison: CnSwe1 residues 795–1084 and CnSwe102 residues 818–1146. Mean pLDDT values are shown for the full-length proteins and kinase-core regions.

**Supplementary Fig. 2.**
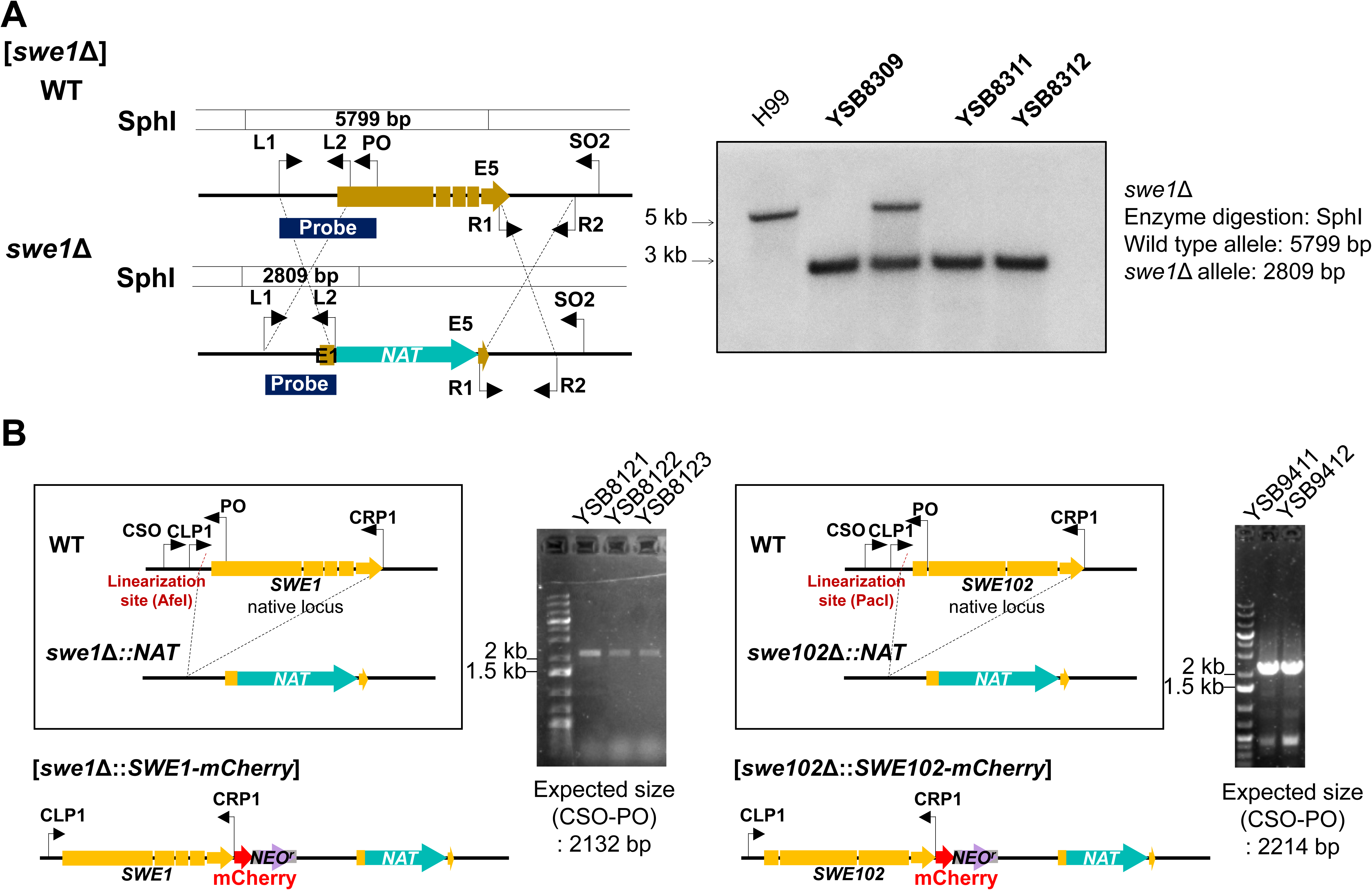
Construction and validation of *swe1*Δ mutant and *SWE1*-mCherry and *SWE102*-mCherry complemented strains. (A) Construction and validation of the *swe1*Δ mutant. The native *SWE1* locus was replaced with the NAT selectable marker by homologous recombination. Correct integration was confirmed by Southern blot analysis after SphI digestion using the indicated probe. The expected band sizes were 5,799 bp for the WT allele and 2,809 bp for the *swe1*Δ allele. (B) Construction and validation of *SWE1*-mCherry and *SWE102*-mCherry complemented strains in the corresponding deletion mutant backgrounds. Schematic diagrams show integration of the mCherry-tagged alleles at the native loci. Correct integration was confirmed by diagnostic PCR using the indicated primer pairs. The expected PCR product sizes were 2,214 bp for *swe1*Δ::*SWE1*-mCherry and 2,132 bp for *swe102*Δ::*SWE102*-mCherry.

**Supplementary Fig. 3.**
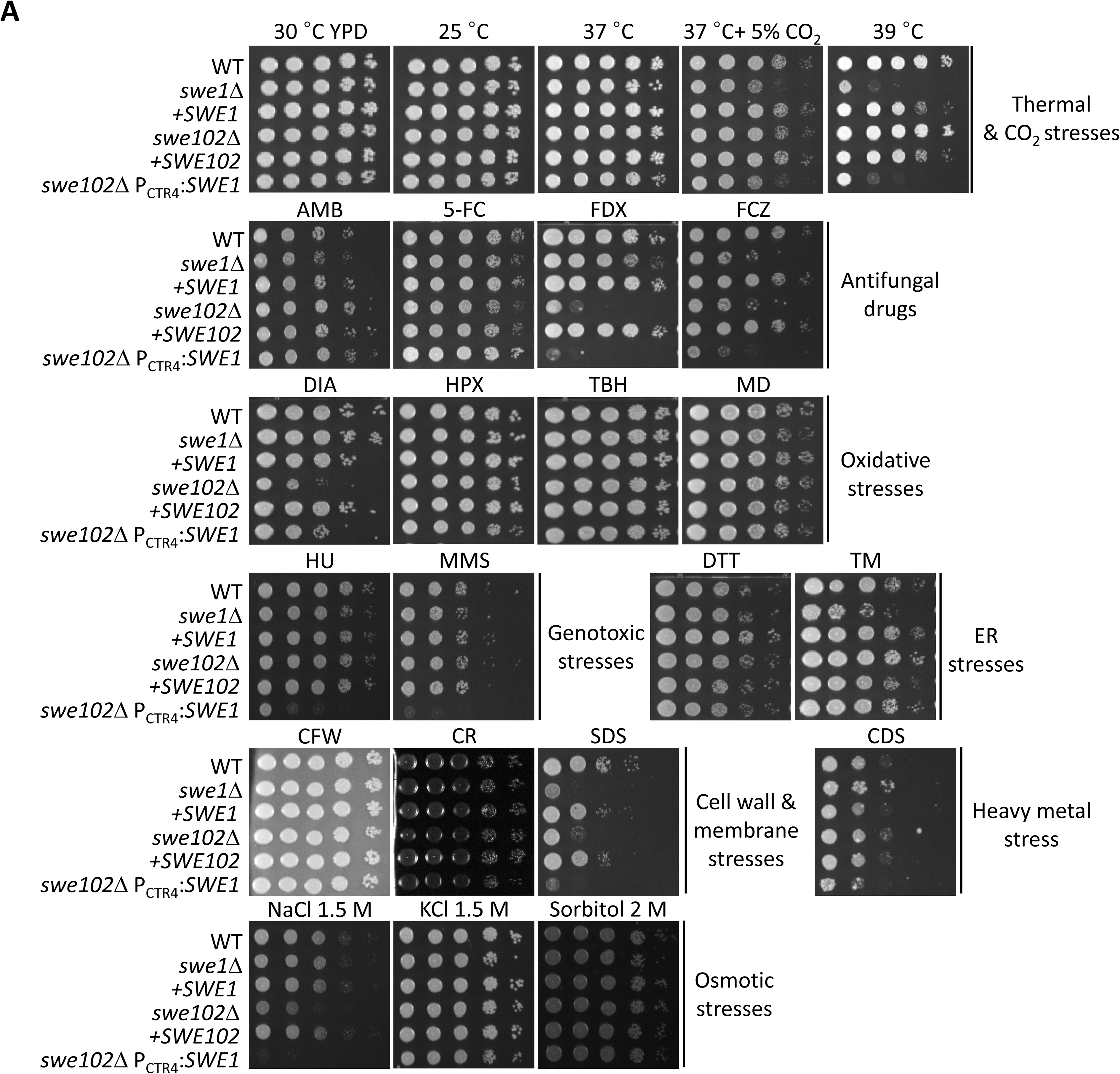
Extended in vitro stress susceptibility profiles of *SWE1* and *SWE102* mutant strains. Serial-dilution growth analysis of WT, *swe1*Δ, *swe102*Δ, *swe1*Δ::*SWE1*-mCherry, *swe102*Δ::*SWE102*-mCherry, and *swe102*Δ *P*CTR4:*SWE1* strains under diverse in vitro stress conditions. Cells were serially diluted 10-fold and spotted onto YPD agar under thermal and CO stress conditions or onto YPD agar supplemented with the indicated stress-inducing agents. Conditions included cell wall and membrane stresses [calcofluor white (CFW; 3 mg/ml), Congo red (CR; 0.8%), sodium dodecyl sulfate (SDS; 0.03%), and cadmium sulfate (CDS; 25 μM)], osmotic stresses [NaCl (1.5 M), KCl (1.5 M), and sorbitol (2 M)], antifungal drugs [amphotericin B (AMB; 1.6 μg/ml), 5-flucytosine (5-FC; 300 μg/ml), fludioxonil (FDX; 1 μg/ml), and fluconazole (FCZ; 10 μg/ml)], oxidative stresses [hydrogen peroxide (HPX; 3 mM), tert-butyl hydroperoxide (TBH; 0.6 mM), menadione (MD; 0.02 mM), and diamide (DIA; 2 mM)], ER stresses [dithiothreitol (DTT; 16 mM) and tunicamycin (TM; 0.3 μg/ml)], and genotoxic stresses [hydroxyurea (HU; 100 mM) and methyl methanesulfonate (MMS; 0.03%)]. Plates were photographed after 3–4 days of incubation.

**Supplementary Fig. 4.**
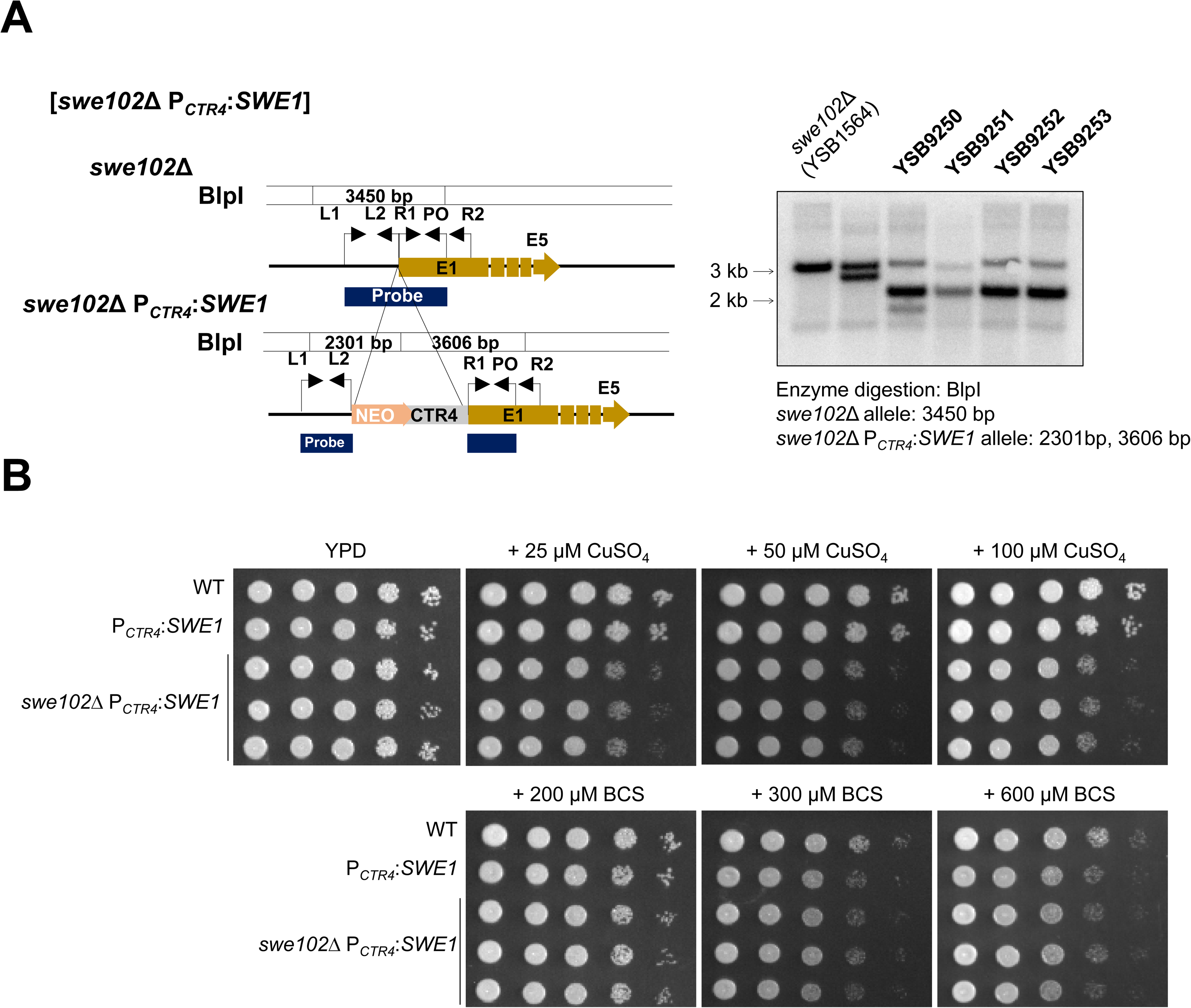
Construction and dose-dependent regulation of the *swe102*Δ P*_CTR4_*:*SWE1* strain. (A) Construction and validation of the *swe102*Δ P*_CTR4_*:*SWE1* strain. Schematic representation of the P*_CTR4_*:*SWE1* promoter-replacement strategy in the *swe102*Δ background is shown. Correct integration was confirmed by Southern blot analysis. Genomic DNA was digested with BlpI and hybridized with the indicated probe. The expected band size for the parental *swe102*Δ allele was 3,450 bp, whereas the correctly integrated *swe102*Δ P*_CTR4_*:*SWE1* allele produced bands of 2,301 bp and 3,606 bp. (B) Serial-dilution growth analysis of WT, P*_CTR4_*:*SWE1*, and *swe102*Δ P*_CTR4_*:*SWE1* strains under copper-repressive and copper-depleted conditions. Cells were spotted onto YPD medium or YPD medium supplemented with CuSO_4_ (25, 50, or 100 μM) or bathocuproinedisulfonic acid (BCS; 200, 300, or 600 μM), as indicated. Plates were photographed after 3–4 days of incubation.

**Supplementary Fig. 5.**
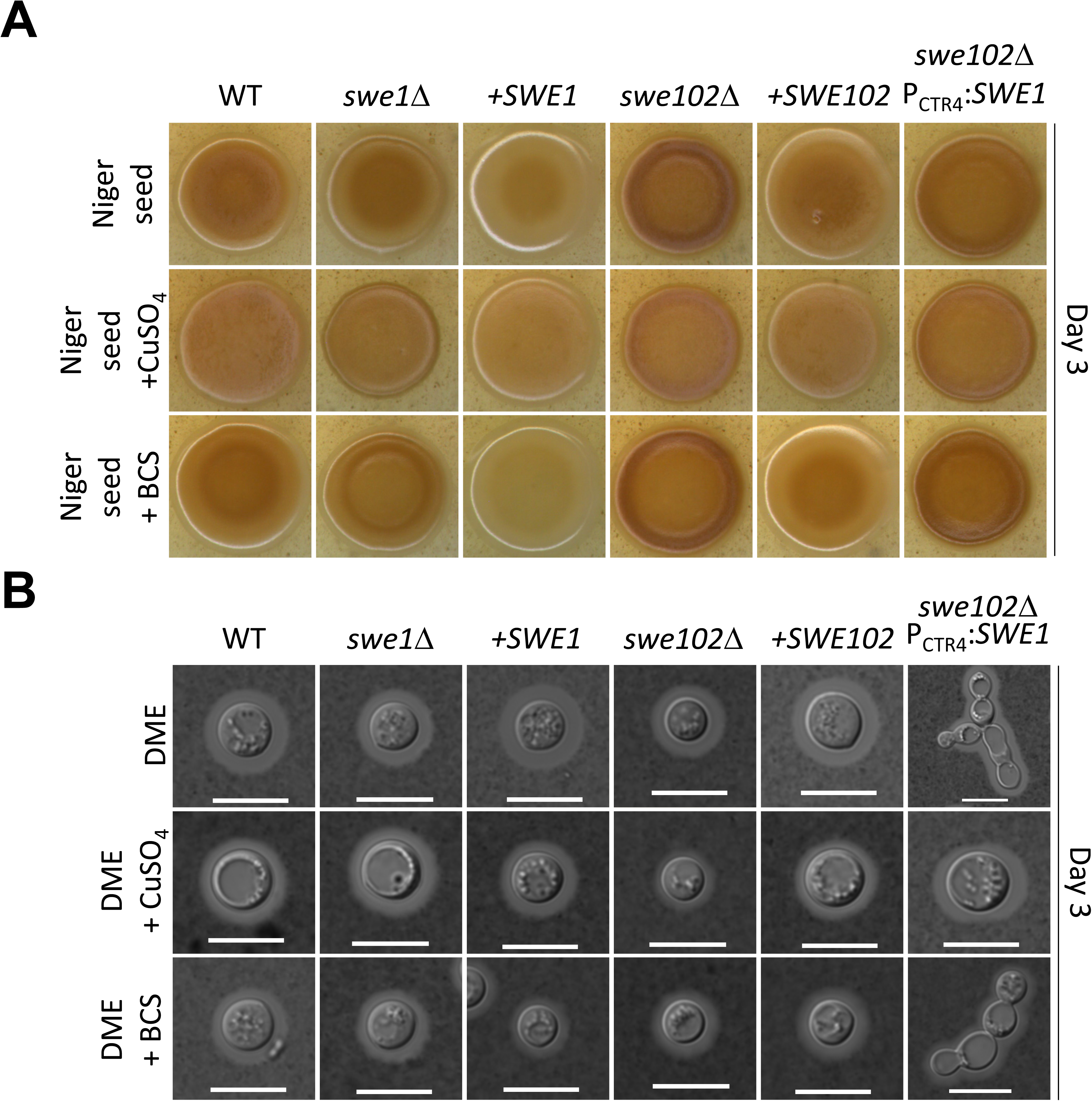
Virulence factor-associated phenotypes of *SWE1*, *SWE102*, and *SWE1*-dosage-regulated strains. Analysis of virulence factor-associated phenotypes in WT, *swe1*Δ, *swe1*Δ::*SWE1*-mCherry, *swe102*Δ, *swe102*Δ::*SWE102*-mCherry, and *swe102*Δ P*_CTR4_*:*SWE1* strains. Cells were spotted onto DME medium to assess capsule-associated phenotypes and onto Niger seed medium to assess melanin production. Media were left untreated or supplemented with CuSO_4_ (25 μM) or BCS (200 μM), as indicated. Plates were photographed after 3 days of incubation.

